# Characterization of extracellular vesicles and synthetic nanoparticles with four orthogonal single-particle analysis platforms

**DOI:** 10.1101/2020.08.04.237156

**Authors:** Tanina Arab, Emily R. Mallick, Yiyao Huang, Liang Dong, Zhaohao Liao, Zezhou Zhao, Olesia Gololobova, Barbara Smith, Norman J. Haughey, Kenneth J. Pienta, Barbara S. Slusher, Patrick M. Tarwater, Juan Pablo Tosar, Angela M. Zivkovic, Wyatt N. Vreeland, Michael E. Paulaitis, Kenneth W. Witwer

**Affiliations:** Department of Molecular and Comparative Pathobiology, Johns Hopkins University School of Medicine, Baltimore, MD, US; Department of Urology, Johns Hopkins University School of Medicine, Baltimore, MD, US; Department of Cell Biology, Johns Hopkins University School of Medicine, Baltimore, MD, US; Department of Neurology, Johns Hopkins University School of Medicine, Baltimore, MD, US; Johns Hopkins Drug Discovery, Johns Hopkins University School of Medicine, Baltimore, MD; Department of Epidemiology, Johns Hopkins University Bloomberg School of Public Health, Baltimore, MD, US; Faculty of Science, Universidad de la República, Montevideo, Uruguay; Functional Genomics Unit, Institut Pasteur de Montevideo, Montevideo, Uruguay; Department of Nutrition, University of California Davis, Davis, CA, US; Bioprocess Measurements Group, National Institute of Standards and Technology, Gaithersburg, MD, US; Center for Nanomedicine at the Wilmer Eye Institute, Johns Hopkins University School of Medicine, Baltimore, MD, US; The Richman Family Precision Medicine Center of Excellence in Alzheimer’s Disease, Johns Hopkins University School of Medicine, Johns Hopkins Medicine and Johns Hopkins Bayview Medical Center, Baltimore, MD, US

**Keywords:** extracellular vesicles, exosomes, microvesicles, ectosomes, nanoparticle tracking analysis, single particle interferometric reflectance imaging sensing, resistive pulse sensing, nanoflow cytometry

## Abstract

We compared four orthogonal technologies for sizing, counting, and phenotyping of extracellular vesicles (EVs) and synthetic particles. The platforms were: single-particle interferometric reflectance imaging sensing (SP-IRIS) with fluorescence, nanoparticle tracking analysis (NTA) with fluorescence, microfluidic resistive pulse sensing (MRPS), and nanoflow cytometry measurement (NFCM). EVs from the human T lymphocyte line H9 (high CD81, low CD63) and the promonocytic line U937 (low CD81, high CD63) were separated from culture conditioned medium (CCM) by differential ultracentrifugation (dUC) or a combination of ultrafiltration (UF) and size exclusion chromatography (SEC) and characterized by transmission electron microscopy (TEM) and Western blot (WB). Mixtures of synthetic particles (silica and polystyrene spheres) with known sizes and/or concentrations were also tested. MRPS and NFCM returned similar particle counts, while NTA detected counts approximately one order of magnitude lower for EVs, but not for synthetic particles. SP-IRIS events could not be used to estimate particle concentrations. For sizing, SP-IRIS, MRPS, and NFCM returned similar size profiles, with smaller sizes predominating (per power law distribution), but with sensitivity typically dropping off below diameters of 60 nm. NTA detected a population of particles with a mode diameter greater than 100 nm. Additionally, SP-IRIS, MRPS, and NFCM were able to identify at least three of four distinct size populations in a mixture of silica or polystyrene nanoparticles. Finally, for tetraspanin phenotyping, the SP-IRIS platform in fluorescence mode was able to detect at least two markers on the same particle, while NFCM detected either CD81 or CD63. Based on the results of this study, we can draw conclusions about existing single-particle analysis capabilities that may be useful for EV biomarker development and mechanistic studies.

## INTRODUCTION

Classification of extracellular vesicles (EVs) into subtypes has been proposed based on size, biogenesis pathway, separation procedure, cellular or tissue origin, and function, among others [1–6]. However, reproducible classification of EV subtypes will require single-particle characterization techniques including phenotyping by surface molecules or molecular signatures [7,8]. In this sense, current knowledge of EV subtypes could be compared with knowledge of immune cells in the 1970s and early 1980s. Around that time, multiplexed flow cytometry capabilities and cell sorting were developed, allowing more precise identification, characterization, and molecular and functional profiling of immune cell subsets [9]. Single-particle technologies for much smaller biological entities will be needed to divide heterogeneous EV populations into well-defined and easily recognized subgroups.

In this study, we evaluated several particle types and single-particle characterization platforms. For input, we used a selection of biological and synthetic particles. EVs were separated from culture medium of H9 T lymphocytic cells and U937 promonocytic cells using several methods. These two cell lines were chosen because they display different levels of the tetraspanins CD63 and CD81. H9 cells have high CD81 and low CD63 levels, while U937 produce little CD81 but abundant CD63. Mixtures of distinct sizes of synthetic silica and traceable polystyrene beads were also tested, not because they mimic EVs or can serve as EV reference materials, but precisely because of their known size and composition, creating a “best case scenario” to assess ability to measure particles. The technology platforms (Text Box 1) were: single-particle interferometric reflectance imaging sensing (SP-IRIS, NanoView) [10–12] with fluorescence, nanoparticle tracking analysis (NTA, ParticleMetrix) [13–15] with fluorescence, microfluidic resistive pulse sensing (MRPS, Spectradyne) [15–17] (which does not have fluorescence capabilities), and nanoflow cytometry measurement (NFCM, NanoFCM) [18,19] with fluorescence.

### Text Box 1: Evaluated Technologies

Single-particle interferometric reflectance imaging sensing (SP-IRIS) captures particles (*e*.*g*. EVs) onto a chip by affinity reagents, usually antibodies, to surface antigens. Particles are imaged by interferometric reflectance for sizing and counting, and fluorescence detection may be done for up to three channels for surface antigens or internal molecules following fixation and permeabilization. Website for the platform we used: https://www.nanoviewbio.com/

Nanoparticle tracking analysis (NTA) is an optical method to track single particles and assign sizes and counts. Measuring Brownian motion allows calculation of a hydrodynamic sphere-equivalent radius of each tracked particle. Additionally, fluorescence filters can be used for detection of particle-associated fluorescence moieties channels. Website for the platform we used: https://www.particle-metrix.de/en/particle-metrix

Microfluidic resistive pulse sensing (MRPS) counts and sizes particles as they pass through a pore between microfluidic chambers. Occlusion of the pore results in a measurable change in electrical signal (defining an event) that is proportional to the volume of the particle. Often, this technique uses different disposable cartridge pore sizes to detect particle populations within specific size ranges. As a non-optical technology, fluorescence detection is not available. Website for the platform we used: https://nanoparticleanalyzer.com/

Nanoflow cytometry measurement (NFCM) is a flow-based technique that detects nano-sized particles by scatter and/or fluorescence. Compared with traditional flow cytometry, a smaller flow channel reduces background signal, and lower system pressure increases dwell time of particles for enhanced signal integration. Website for the platform we used: http://www.nanofcm.com/products/flow-nanoanalyzer

## MATERIALS AND METHODS

*Please see Table 1 for manufacturer, part number, and (where applicable) dilution of reagents*. ***Certain commercial equipment, instruments, and reagents are identified in this paper to foster understanding. Such identification does not imply recommendation or endorsement by the National Institute of Standards and Technology or any other entity, nor does it imply that the materials or equipment identified are necessarily the best available for the purpose***.

**Table 1.**
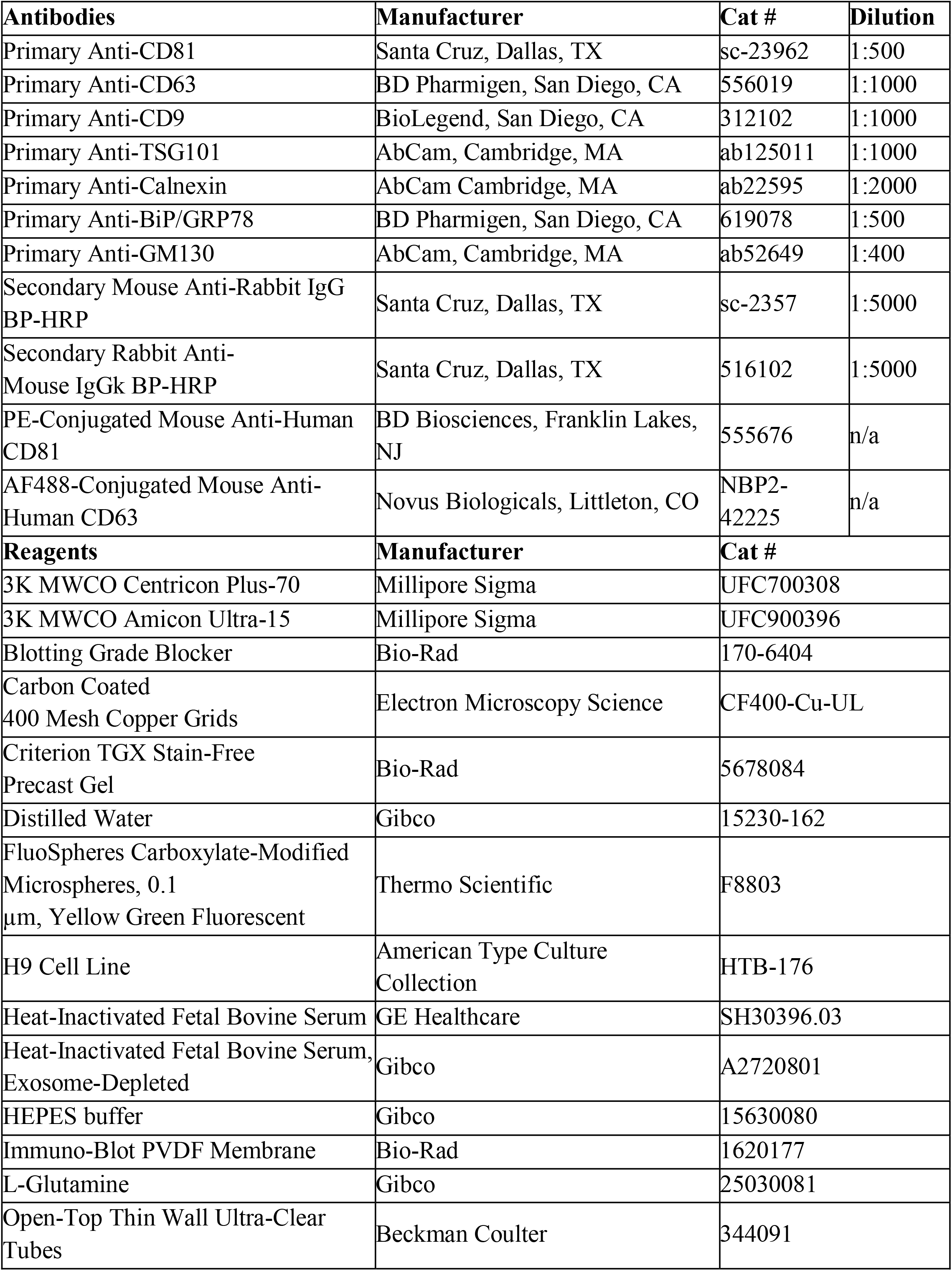
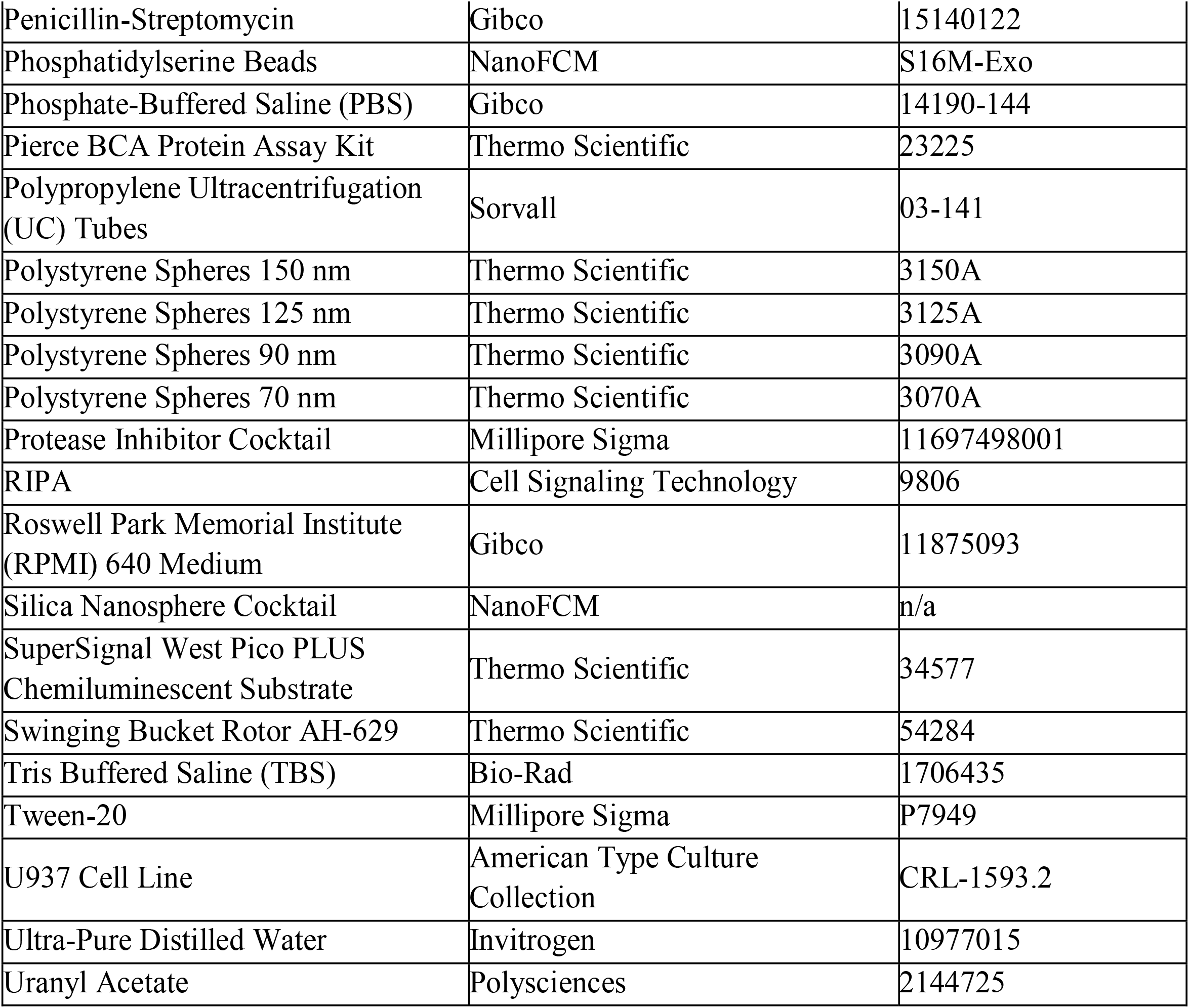

| **Antibodies** | **Manufacturer** | **Cat #** | **Dilution** |
| --- | --- | --- | --- |
| Primary Anti-CD81 | Santa Cruz, Dallas, TX | sc-23962 | 1:500 |
| Primary Anti-CD63 | BD Pharmigen, San Diego, CA | 556019 | 1:1000 |
| Primary Anti-CD9 | BioLegend, San Diego, CA | 312102 | 1:1000 |
| Primary Anti-TSG101 | AbCam, Cambridge, MA | ab125011 | 1:1000 |
| Primary Anti-Calnexin | AbCam Cambridge, MA | ab22595 | 1:2000 |
| Primary Anti-BiP/GRP78 | BD Pharmigen, San Diego, CA | 619078 | 1:500 |
| Primary Anti-GM130 | AbCam, Cambridge, MA | ab52649 | 1:400 |
| Secondary Mouse Anti-Rabbit IgG BP-HRP | Santa Cruz, Dallas, TX | sc-2357 | 1:5000 |
| Secondary Rabbit Anti-Mouse IgGk BP-HRP | Santa Cruz, Dallas, TX | 516102 | 1:5000 |
| PE-Conjugated Mouse Anti-Human CD81 | BD Biosciences, Franklin Lakes, NJ | 555676 | n/a |
| AF488-Conjugated Mouse Anti-Human CD63 | Novus Biologicals, Littleton, CO | NBP2-42225 | n/a |
| **Reagents** | **Manufacturer** | **Cat #** |  |
| 3K MWCO Centricon Plus-70 | Millipore Sigma | UFC700308 |  |
| 3K MWCO Amicon Ultra-15 | Millipore Sigma | UFC900396 |  |
| Blotting Grade Blocker | Bio-Rad | 170-6404 |  |
| Carbon Coated 400 Mesh Copper Grids | Electron Microscopy Science | CF400-Cu-UL |  |
| Criterion TGX Stain-Free Precast Gel | Bio-Rad | 5678084 |  |
| Distilled Water | Gibco | 15230-162 |  |
| FluoSpheres Carboxylate-Modified Microspheres, 0.1 μm, Yellow Green Fluorescent | Thermo Scientific | F8803 | |
| H9 Cell Line | American Type Culture Collection | HTB-176 |  |
| Heat-Inactivated Fetal Bovine Serum | GE Healthcare | SH30396.03 |  |
| Heat-Inactivated Fetal Bovine Serum, Exosome-Depleted | Gibco | A2720801 |  |
| HEPES buffer | Gibco | 15630080 |  |
| Immuno-Blot PVDF Membrane | Bio-Rad | 1620177 |  |
| L-Glutamine | Gibco | 25030081 |  |
| Open-Top Thin Wall Ultra-Clear Tubes | Beckman Coulter | 344091 |  |

|  |  |  |
| --- | --- | --- |
| Penicillin-Streptomycin | Gibco | 15140122 |
| Phosphatidylserine Beads | NanoFCM | S16M-Exo |
| Phosphate-Buffered Saline (PBS) | Gibco | 14190-144 |
| Pierce BCA Protein Assay Kit | Thermo Scientific | 23225 |
| Polypropylene Ultracentrifugation (UC) Tubes | Sorvall | 03-141 |
| Polystyrene Spheres 150 nm | Thermo Scientific | 3150A |
| Polystyrene Spheres 125 nm | Thermo Scientific | 3125A |
| Polystyrene Spheres 90 nm | Thermo Scientific | 3090A |
| Polystyrene Spheres 70 nm | Thermo Scientific | 3070A |
| Protease Inhibitor Cocktail | Millipore Sigma | 11697498001 |
| RIPA | Cell Signaling Technology | 9806 |
| Roswell Park Memorial Institute (RPMI) 640 Medium | Gibco | 11875093 |
| Silica Nanosphere Cocktail | NanoFCM | n/a |
| SuperSignal West Pico PLUS Chemiluminescent Substrate | Thermo Scientific | 34577 |
| Swinging Bucket Rotor AH-629 | Thermo Scientific | 54284 |
| Tris Buffered Saline (TBS) | Bio-Rad | 1706435 |
| Tween-20 | Millipore Sigma | P7949 |
| U937 Cell Line | American Type Culture Collection | CRL-1593.2 |
| Ultra-Pure Distilled Water | Invitrogen | 10977015 |
| Uranyl Acetate | Polysciences | 2144725 |

### Particle preparation

Human cells lines H9 (T lymphocytic) and U937 (pro-monocytic) were obtained from the American Type Culture Collection (ATCC). Cells were maintained in Roswell Park Memorial Institute (RPMI) 1640 Medium supplemented with either replete or EV-depleted 10% heat-inactivated fetal bovine serum, with 1% HEPES buffer, 1% Penicillin-Streptomycin, and 1% L-Glutamine. Cells were cultured at 37 °C in 5% CO_2_. Silica spheres (SS, NanoFCM, Nottingham, England) were a premixed combination of diameters 68 nm, 91 nm, 113 nm, and 151 nm. Individual polystyrene spheres (PS, Thermo Fisher) of diameters 70 nm, 90 nm, 125 nm, and 150 nm were purchased. Nominally equal concentrations (1 × 10^12^ particles/mL) of beads were mixed.

### Size-exclusion chromatography (SEC)

60 mL of culture-conditioned medium (CCM) from each cell line was centrifuged at 1,000 × g for 5 minutes at 4 °C to remove cells and cellular debris. 3 kDa molecular weight cut off (MWCO) Centricon Plus-70 centrifugal filters (Millipore Sigma) were used to concentrate the initial volume to 1.5 mL. Size exclusion chromatography (SEC) was done with qEV Automated Fraction Collectors (AFC; Izon Science, Cambridge, MA) and qEV original 70 nm columns (Izon Science, Cambridge, MA). Columns were left at room temperature for 30 minutes and washed with phosphate-buffered saline (PBS). 0.5 mL of concentrated CCM was loaded onto each of three columns, and 0.5 mL fractions were collected by adding additional PBS to the column. EV-enriched fractions (SEC; fractions 7-9) were pooled altogether from the three columns used for each sample and further concentrated using 3 kDa MWCO Amicon Ultra-15 Centrifugal Filters to a final volume of 1 mL. 50-µL aliquots were stored at −20 °C for downstream assays.

### Differential ultracentrifugation (dUC)

60 mL of CCM from each cell line was centrifuged at 1,000 × g for 5 minutes at 4 °C to remove cells and cellular debris and 2,000 × g for 10 minutes at 4 °C to remove additional debris. The supernatant was transferred to polypropylene thin-wall ultracentrifugation (UC) tubes and centrifuged at 10,000 × g for 30 minutes at 4 °C using a swinging bucket rotor (Thermo Scientific rotor model AH-629, k-factor 242, acceleration and deceleration settings of 9) to pellet large EVs. Supernatant was transferred into new polypropylene thin wall UC tubes and centrifuged at 100,000 × g for 70 minutes at 4 °C using the same swinging bucket rotor. The 100K pellets containing small EVs were resuspended in 1 mL of PBS, vigorously vortexed, and placed on ice for 20 minutes. 50-µL aliquots were stored at −20 °C for downstream assays.

### Transmission electron microscopy (TEM)

10 µL freshly thawed aliquots were adsorbed to glow-discharged carbon-coated 400 mesh copper grids by flotation for 2 minutes. Grids were quickly blotted and rinsed by flotation on 3 drops (1 minute each) of 1× Tris-buffered saline. Grids were negatively stained in 2 consecutive drops of 1% uranyl acetate (UAT) with tylose (1% UAT in deionized water (dIH_2_O), double filtered through a 0.22 µm filter), blotted, then quickly aspirated to cover the sample with a thin layer of stain. Grids were imaged on a Hitachi 7600 TEM operating at 80 kV with an AMT XR80 CCD (8 megapixel). SS and PS were absorbed to grids as above, but with initial flotation for 5 minutes and imaging on a Phillips CM-120 TEM operating at 80 kV with an AMT XR80 CCD (8 megapixel).

### Western blot (WB)

H9 and U937 cell pellets and isolated EVs were lysed in 1× radioimmunoprecipitation assay buffer (RIPA) supplemented with protease inhibitor cocktail. Protein quantification of cell and EV lysates was done using a bicinchoninic acid assay (BCA) (Pierce BCA Protein Assay Kit). 5 µg of lysates were resolved using a 4% to 15% Criterion TGX Stain-Free Precast gel, then transferred onto an Immuno-Blot PVDF membrane. Blots were probed using primary antibodies in PBS-T and 5% Blotting Grade Blocker. Primary antibodies were against CD81, CD63, CD9, TSG101, calnexin, BiP/GRP78, and GM130. Secondary antibodies were rabbit anti-mouse IgGk BP-HRP and mouse anti-rabbit IgGk BP-HRP. SuperSignal West Pico PLUS Chemiluminescent Substrate was used for detection and blots were visualized with an iBright Western Blot (Thermo Fisher, Waltham, MA) imaging system.

### Single particle interferometric reflectance imaging (SP-IRIS)

Measurements were performed largely as described previously [20,21]. 35 µL of H9 and U937 EVs isolated by SEC or dUC were diluted 1:1 in incubation buffer (IB) and incubated at room temperature on ExoView R100 (NanoView Biosciences, Brighton, MA) chips printed with anti-human CD81 (JS-81), anti-human CD63 (H5C6), anti-human CD9 (HI9a), and anti-mouse IgG1 (MOPC-21).

After incubation for 16 hours, chips were washed with IB 4 times for 3 minutes each under gentle horizontal agitation at 500 rpm. Chips were then incubated for 1 hour at room temperature with a fluorescent antibody cocktail of anti-human CD81 (JS-81, CF555), anti-human CD63 (H5C6, CF647), and anti-human CD9 (HI9a, CF488A) at a dilution of 1:1200 (v:v) in a 1:1 (v:v) mixture of IB and blocking buffer. The buffer was then exchanged to IB only, followed by 1 wash with IB, 3 washes with wash buffer, and 1 wash with rinse buffer (3 minutes each at 500 rpm). Chips were immersed twice in rinse buffer for approximately 5 seconds each and removed at a 45-degree angle to allow the liquid to vacate the chip. All reagents and antibodies were supplied by NanoView Biosciences (Brighton, MA, Cat #EV-TETRA-C). Both SS and PS were diluted in dIH_2_O to load 10,000 particles, nominally, per antibody capture spot on the ExoView chips. 35 µL of diluted spheres were incubated on ExoView chips and allowed to fully dry. All chips were imaged in the ExoView scanner (NanoView Biosciences, Brighton, MA) by interferometric reflectance imaging and fluorescent detection. Data were analyzed using NanoViewer 2.8.10 Software (NanoView Biosciences). Fluorescent cutoffs were as follows: CF555 channel 230, CF488 channel 475, CF647 channel 250 (biological particles) and CF555 channel 675, CF488 channel 600, and CF647 channel 375 (SS and PS).

### Nanoparticle tracking analysis (NTA)

ZetaView QUATT-NTA Nanoparticle Tracking-Video Microscope PMX-420 and BASIC NTA-Nanoparticle Tracking Video Microscope PMX-120 (Particle Metrix, Inning am Ammersee, Germany) instruments were used for particle quantification in both scatter and fluorescence (488 nm) modes. Calibration beads and biological samples were diluted in distilled water and PBS, respectively, to a final volume of 1 mL. Calibration was done for both scatter and fluorescence measurements. For scatter-mode calibration, 100 nm PS beads were diluted 1:250,000 (v:v). Capture settings were: sensitivity 65, shutter 100, minimum trace length 10. Cell temperature was maintained at 25 °C for all measurements. For fluorescence calibration, 488 nm yellow-green FluoSpheres were diluted 1:250,000 (v:v), and both scatter and fluorescence were measured. Scatter was recorded as above, and fluorescence was measured at sensitivity 80, shutter 100, and minimum trace length 15. To measure SS and PS mixtures and individual size populations of PS, samples were diluted such that at least 200 particles could be counted per frame. Technical triplicates were measured for each sample. A washing step was done between each measurement using dIH_2_O. For H9 and U937 EVs separated by SEC or dUC, one cycle was performed by scanning 11 cell positions. Capture was done at medium video setting, corresponding to 30 frames per position. ZetaView software 8.5.10 was used to analyze the recorded videos with the following settings: minimum brightness 30, maximum brightness 255, minimum area 10, and maximum area 1000. Since subpopulations of particles might also be identified based on signal intensity, we used manual and population distribution gates in the ZetaView software to assess this possibility for SS and PS mixtures. PE-conjugated mouse anti-human CD81 and AF488-conjugated mouse anti-human CD63 were used for fluorescence detection of EVs. Antibodies were mixed 1:9 (v:v) with PBS, incubated 2 hours at room temperature, and diluted to a final volume of 1 mL. Supplementary Table 2 lists all antibodies tested with this platform.

### Microfluidic resistive pulse sensing (MRPS)

Microfluidics resistive pulse sensing measurements were conducted using the nCS1 instrument (Spectradyne, Torrance, CA) as described previously [20]. For biological particles, sample volumes of a few µL of H9 and U937 EVs isolated by SEC or dUC were diluted with an equal volume of 1% polysorbate 20 (Tween 20) in 1× PBS (PBST) and further diluted as indicated with 1× PBS, and loaded onto polydimethylsiloxane cartridges (diameter range 65 nm to 400 nm). A different cartridge was used for each sample and replicate. Approximately 5 µL of the diluted sample was used and about 25,000 events were recorded for each analyte. For synthetic nanoparticles, SS and PS were diluted 100-fold by volume in dIH_2_O, then 10-fold by volume with equal volumes of PBST and the remainder with 1× PBS and loaded onto TS-400 polydimethylsiloxane cartridges. Approximately 3,000 events were obtained for each SS and PS repeat. All acquired results were analyzed using the nCS1 Data Analyzer (Spectradyne, Torrance, CA). For all samples, user-defined filtering was applied by defining 2D polygonal boundaries based on transition time and diameter to exclude false positive signals, similar to gating commonly used in analyzing flow cytometry data. Effects of Tween 20 on EV integrity or counts were assessed by diluting samples to a final concentration of Tween 20 (in PBS) ranging from 0.1% to 0.9%.

### Nano-flow cytometry measurement (NFCM)

The nFCM flow nano-analyzer was used to measure concentration and size of particles following the manufacturer’s instructions and as described previously [22]. Briefly, two single photon-counting avalanche photodiodes (APDs) were used for the simultaneous detection of side scatter (SSC) and fluorescence of individual particles. The instrument was calibrated separately for concentration and size using 200 nm PE- and AF488 fluorophore-conjugated PS beads and a Silica Nanosphere Cocktail, respectively. 20 μL of each EV preparation was incubated with 20 μL PE-conjugated CD81 and 5 μL AF488-conjugated CD63 antibodies at 37 °C for 30 minutes. After incubation, the mixture was washed twice with PBS and centrifuged at 110,000 × g for 70 min at 4 °C (TH-641 rotor, k-factor 114, Thermo Fisher, using thin-wall polypropylene tubes with 13.2 ml capacity and acceleration and deceleration settings of 9). The pellet was resuspended in 50 μL PBS. Events were recorded for 1 minute. Using the calibration curve, the flow rate and side scattering intensity were converted into corresponding particle concentrations and size.

### Dynamic light scattering (DLS)

To check the nominal size values of PS beads, particle diameter was measured by dynamic light scattering using a Malvern Zetasizer Nano-ZS90. Each suspension was diluted 10 × in ultrapure water, and measurements were carried out in triplicate at 25 °C. A single peak was observed for each individual run.

### Data and methods availability

We have submitted all relevant data of our experiments to the EV-TRACK knowledgebase (EV-TRACK ID: EV200090) [23]. Reporting for NFCM was submitted to FlowRepository as ID:FR-FCM-Z2U3 [24]. Any and all data are available on reasonable request.

## RESULTS

### Production, separation, and characterization of input materials

Supernatants were collected from cultured human cell lines: H9 (T-lymphocytic) and U937 (pro-monocytic). Our goal was to obtain EV-enriched or -depleted biological material from cells with different tetraspanin expression. EVs were partially separated by size exclusion chromatography and ultrafiltration or differential ultracentrifugation (Figure 1A). Marker expression and morphology were assessed by WB (Figure 1B and Supplementary Figure 1) and TEM. WB revealed characteristic cellular CD63 and CD81 expression patterns, with CD81 above the limit of detection only for H9 and CD63 predominating for U937 (Figure 1B). CD81 was apparently enriched in EV fractions from H9, while CD63 appeared to be present, but not enriched, in EVs from U937, suggesting relatively inefficient release. Please note, however, that protein amount was used to normalize WB input, so per-particle content cannot be easily compared across sample types, and see also additional blots in Supplementary Figure 1B-D. Calnexin was detected in cell lysates, with little or no signal in EV fractions (Figure 1B). For EVs concentrated and separated by each method, TEM showed heterogeneous populations (particles ranging from approximately 50 to approximately 500 nm in diameter) including EVs with the typical “cup-shaped” fixation artifact (Figure 1C and Supplementary Figure 2). UC pellets displayed higher background and apparent non-EV particles than SEC EV fractions, possibly consistent with proteinaceous material that that elutes in later, relatively EV-depleted fractions of SEC (Figure 1C and Supplementary Figure 2).

**Figure 1:**
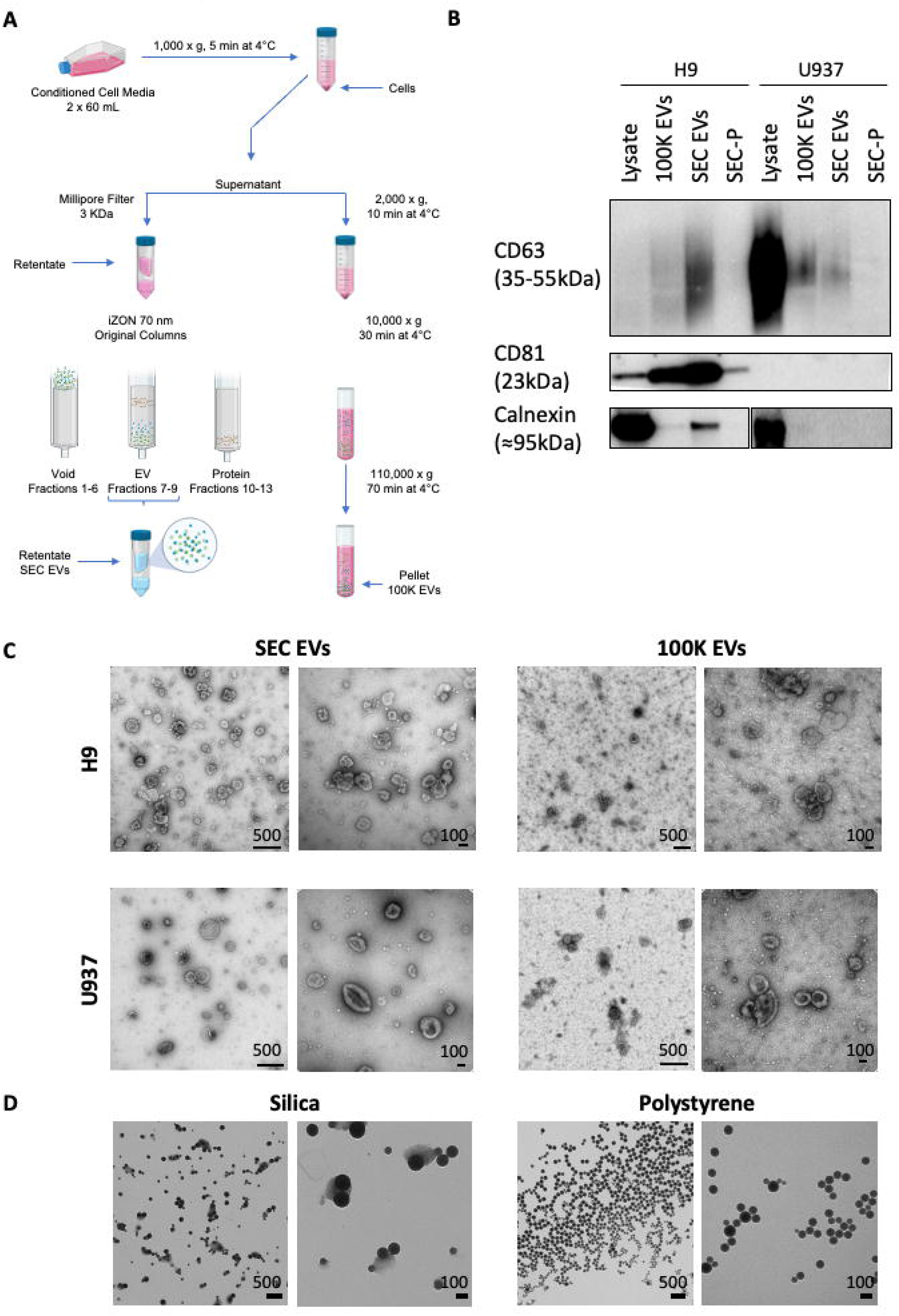
Methodology and EV separation. (A) EVs were separated from H9 and U937 culture-conditioned media by a combination of ultrafiltration and size exclusion chromatography (SEC EVs) or by differential ultracentrifugation (100K EVs). (B) Immunoblots of cell lysates from H9 and U937, EVs separated by ultracentrifugation (100K EVs) and SEC (SEC EVs), and later fractions of SEC (enriched for protein; SEC-P). Antibodies are specified in Table 1; see also Supplementary Figure 1. (C) Electron micrograph of SEC EVs and 100K EVs from both cell lines. As indicated for each subpanel, leftmost scale bars represent 500 nm at magnification 40,000×; rightmost scale bars are 100 nm at magnification 100,000×. (D) EM of SS and PS. Leftmost scale bars are 500 nm at magnification 17,500×; rightmost scale bars are 100 nm at magnification 65,000×.

Silica spheres (SS) and polystyrene spheres (PS) of known size were obtained from commercial sources. These artificial nanoparticles were measured here not as reference materials for EV studies, but simply because they have known diameters and composition, along with higher refractive indices (RIs) than EVs. Note that such particles can be used as reference materials in EV studies if the RI of the material is accounted for, for example with several available software packages [25–28]. For PS, we used National Institute of Standards and Technology (NIST)-traceable size standards. These beads are among the most commonly used size calibrants for materials in their size ranges and are compared with a known standard maintained by NIST. A Certificate of Calibration and Traceability allows labs to show compliance with various ISO and GMP standards and regulations. Additionally, uncertainty of measurement is indicated on a certificate of analysis for each bead lot. We nevertheless confirmed size and purity of SS and PS mixtures using TEM (Figure 1D). Beads corresponding to all four sizes in each mixture were clearly present on the grids, with little or no contaminating material. Bead diameters as measured by TEM were consistent with the nominal diameters and data sheet specifications (Supplementary Table 1, n=at least 30 per population over 4 TEM frames). We also measured individual PS bead populations by dynamic light scattering (DLS), a method best suited for measurement of monodisperse populations (three bead preparations each, measured thrice each). Results showed a single peak for each individual run and polydispersity indices consistent with monodispersity (Supplementary Table 1).

### Artificial nanoparticle sizing

Mixed silica spheres (SS) with nominal diameters of 68 nm, 91 nm, 113 nm, and 151 nm were measured with the four platforms. SP-IRIS identified four distinct populations with diameter modes around 75 nm, 100 nm, 120 nm, and 150 nm (Figure 2A). Since the SP-IRIS technology uses affinity to capture particles, particle mixtures were dried onto the SP-IRIS chips before imaging. NTA detected a broad population distribution with a mode around 105 nm diameter (Figure 2B). MRPS resolved four distinct peaks for each individual chip, but this distinction was masked somewhat by averaging all results (Figure 2C; see inset for an example of an individual reading and also Supplementary Figure 3). NFCM resolved four populations with distinct peaks at diameters of approximately 66 nm, 85 nm, 112 nm, and 154 nm (Figure 2D). Polystyrene spheres (PS) with nominal diameters 70 nm, 90 nm, 125 nm, and 150 nm were mixed to a nominal concentration of 1×10^12^ particles/mL. SP-IRIS detected four distinct peaks around 80 nm, 110 nm, 140 nm, and 170 nm (Figure 2E). NTA returned a broad population distribution centered around 105 nm (Figure 2F). MRPS identified distinct peaks at diameters 71 nm, 92 nm, 123 nm, and 150 nm (Figure 2G). Nano-flow showed four populations around 85 nm, 120 nm, 170 nm, and 225 nm in diameter, as well as a possible smaller population around 60 nm (Figure 2H). We also measured several dilutions of SS and PS particles using MRPS and NFCM, with results similar to those described above. Raw and dilution-corrected data are presented in Supplementary Figure 4.

**Figure 2:**
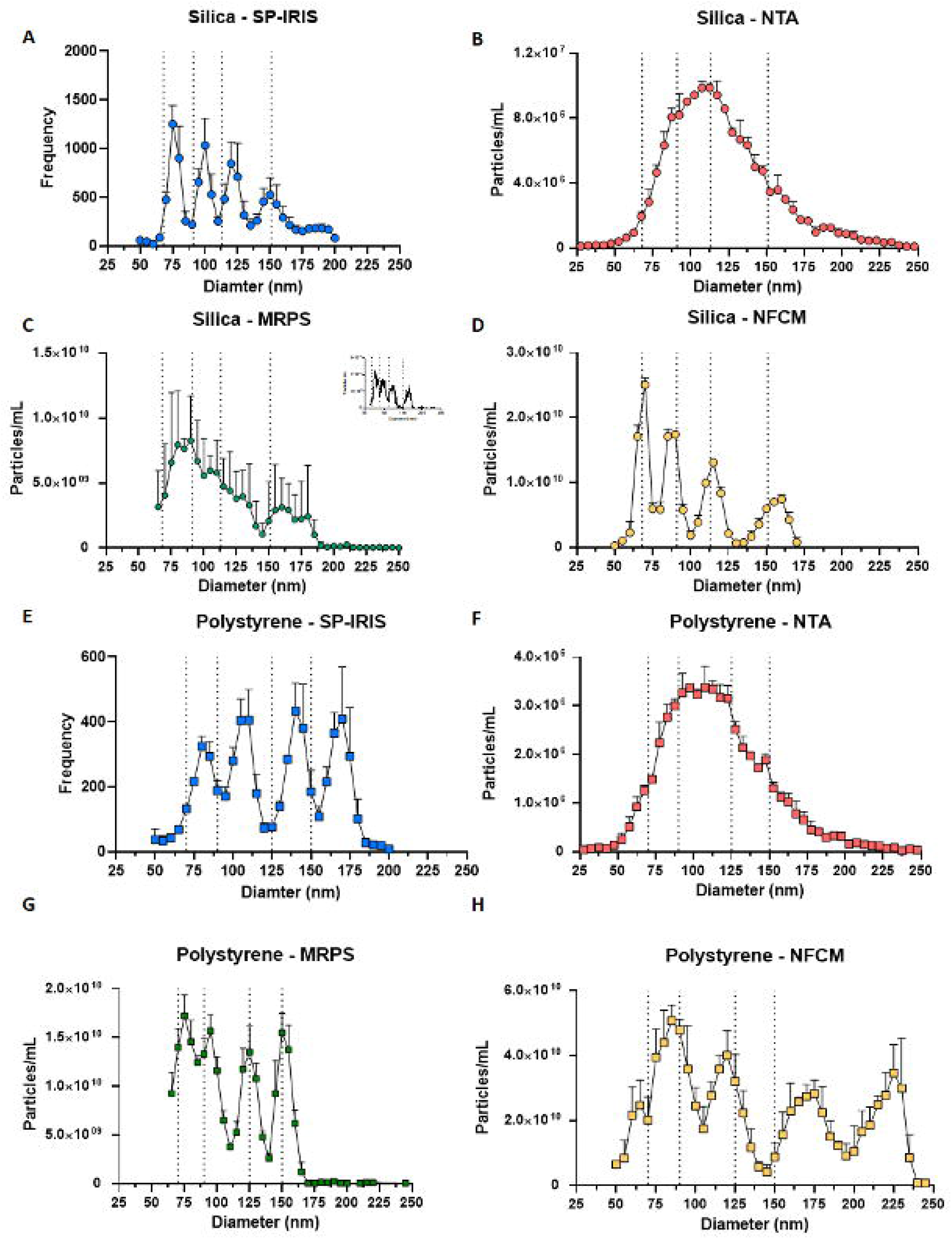
SS and PS size distribution. Size distributions for SS (n=3) with standard deviation for (A) SP-IRIS, (B) NTA, (C) MRPS, and (D) NFCM. Nominal SS diameters are indicated by vertical dotted lines: 68 nm, 91 nm, 113 nm, and 151 nm. Size distributions for PS (n=3; with SD) for (E) SP-IRIS, (F) NTA, (G) MRPS, and (H) NFCM. Nominal PS diameters are indicated by vertical dotted lines: 70 nm, 90 nm, 125 nm, and 150 nm. Inset in Figure 2C shows a single MRPS measurement of the size distribution; see also Supplementary Figure 3 for individual readings.

Because the NTA platform did not appear to resolve different populations, we also assessed individual PS bead sizes and also tried to use intensities to resolve individual bead populations. Individually, the four bead sizes returned measurements (arithmetic mean +/-SD) of 109.0 nm +/-0.4 nm (70 nm PS), 105.2 nm +/-0.3 nm (90 nm PS), 126.5 nm +/-0.4 nm (125 nm PS), and 148.7 nm +/-3.6 nm (150 nm PS). Mixed beads again produced a single broad peak averaging 124.2 nm. Using the NTA software to assign gates based on intensity, we assessed the possibility that individual bead sizes could be resolved. For PS beads, the most intense signals skewed slightly towards larger returned sizes (Supplementary Figure 5A); however, there was no apparent size or distribution difference between medium- and low-intensity populations (Supplementary Figure 5A,B). Similar results were obtained for SS beads (Supplementary Figure 5C). Gating on intensity might, however, be a useful tool in some settings.

### Counting of synthetic nanoparticles

In addition to particle size, we also assessed counts. For SP-IRIS, a mean of around 3000 SS particles were detected per printed antibody spot (Figure 3A), with no overall differences between groups of antibody spots (*i*.*e*., three spots per chip each of three tetraspanins and an isotype control; note that no differences would be expected, since particles were dried onto the chips). However, per-spot events ranged considerably from <2000 SS particles per spot to >4500 SS particles per spot (Figure 3A). SP-IRIS performed similarly for PS. There were no differences between spots printed with different antibodies, with a mean of around 1400 events/antibody spot (Figure 3B), but events per spot ranged from <1000 PS particles/spot to 3000 PS particles/spot. Based on the nominal PS bead concentration and the surface area of the chips, 10,000 particles per spot would have been expected (Figure 3B, dotted line); however, it is possible that beads may not have dried evenly, for example, if they were relatively repelled by antibody-printed surfaces. Following SP-IRIS measurements, chips were probed with three fluorescently labeled antibodies (anti-CD81, anti-CD63, and anti-CD9) to assess background binding. Background binding was negligible for both SS and PS (Supplementary Figure 6A and B, respectively). Some outliers were observed for CD9 (SS) or CD63 (PS); however, none exceeded 1000 events. Particle concentrations were also measured by NTA, MRPS, and NFCM. For SS (Figure 3C), MRPS estimated a concentration approximately one log higher than NTA (5.1×10^11^ particles/mL *vs*. 5.4×10^10^ particles/mL, respectively), with NFCM in the middle (1.7×10^11^ particles/mL). For PS, all three methods were in close agreement (Figure 3D). The difference for SS, but not for PS, likely reflects the sensitivity of the optical techniques to the refractive index of the particles being counted. Furthermore, the measured concentration was very close to the nominal PS concentration of 1×10^12^ particles/mL (Figure 3D, dotted line).

**Figure 3:**
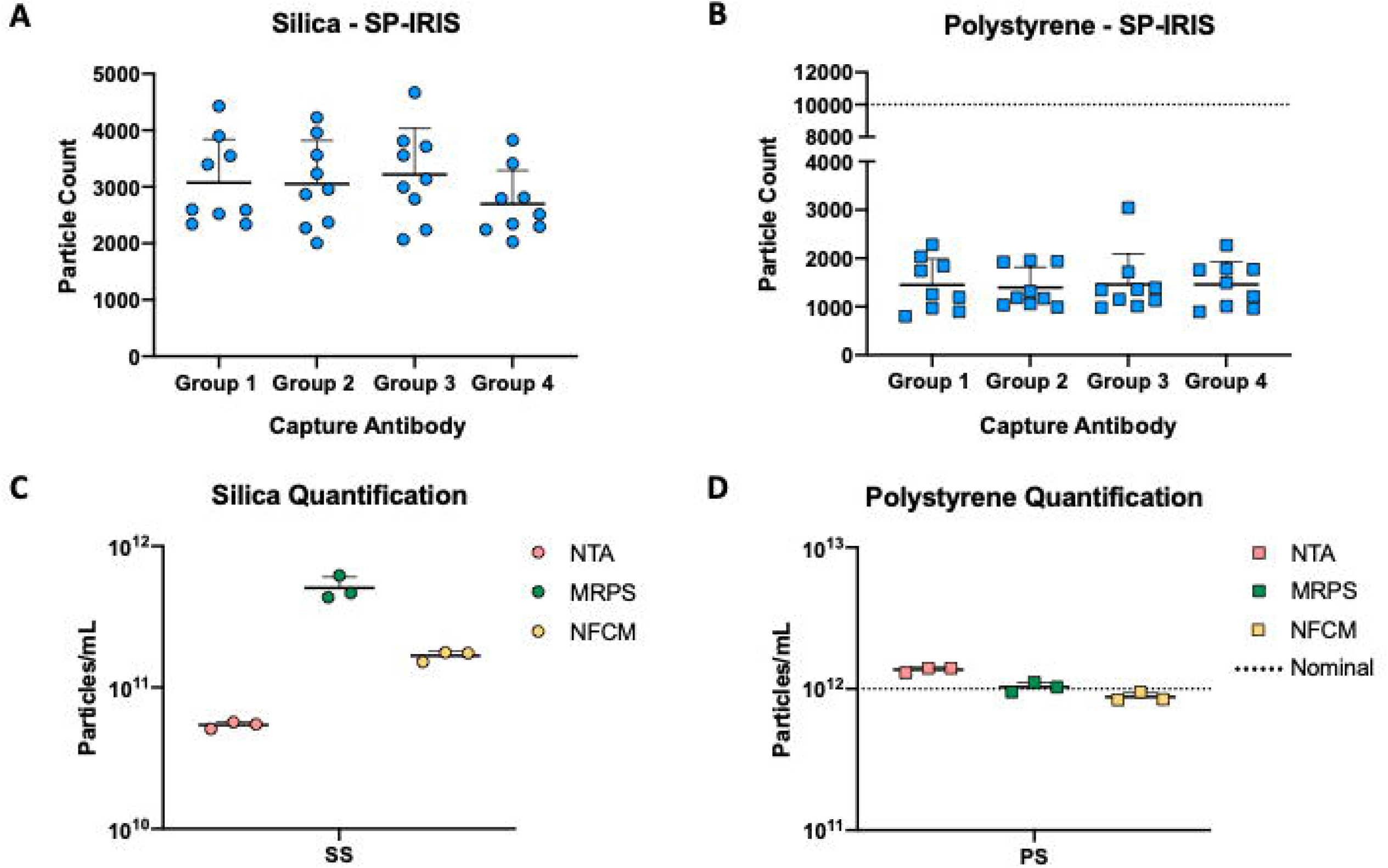
SS and PS quantification. (A) SP-IRIS label-free capture for SS and PS using four capture spots (n=3 per group; mean particle count per spot with SD). B) SS quantification (n=3; mean particles/mL with SD). (C) PS quantification (n=3; mean particles/mL with SD). In panels B and D, nominal PS concentration is indicated by a horizontal dotted line (1.0×10^12^ particles/mL).

### Biological particle sizing

EV preparations from H9 and U937 cell supernatants enriched by ultrafiltration and SEC (SEC EVs) or by differential ultracentrifugation (100K EVs) were next measured using each platform. For H9-derived materials, SP-IRIS returned an almost identical size distribution profile for both EV enrichment methods (Figure 4A). In contrast, NTA, MRPS, and NFCM returned data indicating particles at smaller diameters for the 100K EVs compared with the SEC EVs, with roughly similar particle size distributions (Figure 4B-D). However, substantial variation between replicates might limit the conclusions that can be drawn from this observation; we also expect that the polydisperse nature of EVs will naturally lead to greater CVs. For U937-derived materials, SP-IRIS and NTA (Figure 4E,F) detected more particles at smaller diameters from the 100K EVs compared with the SEC-EVs, again with roughly similar particle size distribution. MRPS produced equivalent particle size distribution and particle number between the two enrichment techniques (Figure 4G). In contrast, NFCM detected a higher particle count of smaller particle diameters from the SEC EVs than the 100K EVs, with the particle size distributions significantly different (Figure 4H). Please see Supplementary Figure 7 for plots drawn without error bars for clarity. Again, variability between replicates limits conclusions. Overall, the results are broadly consistent with the reported power-law size distribution of EVs [29,30] and the expectation that UC pellets may contain non-EV extracellular particles (EPs) around the same size as EVs [1].

**Figure 4:**
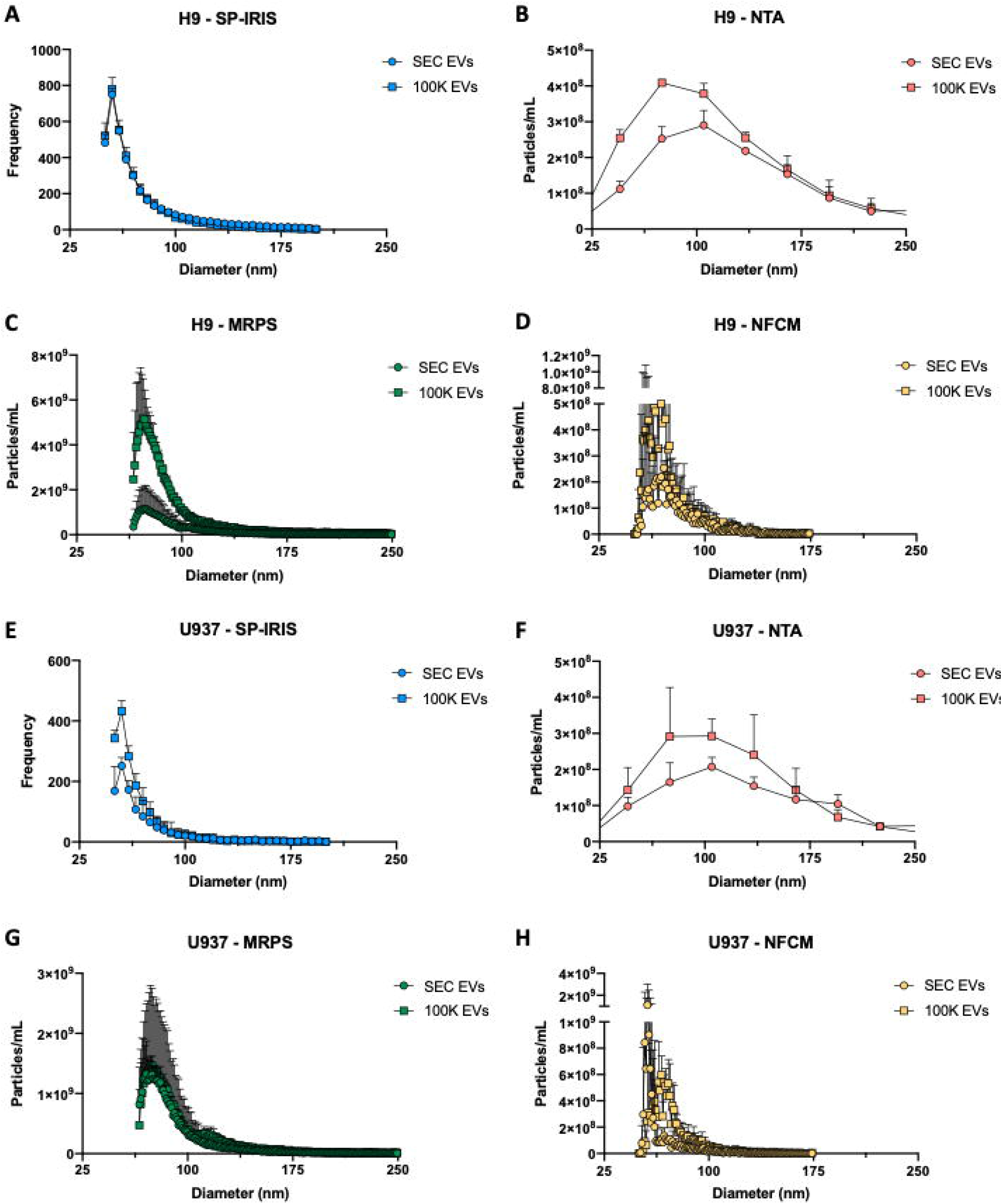
H9 and U937 particle size distribution. Diameters of particles for H9 SEC EVs and 100K EVs (n=3 per group, with standard deviation) for (A) SP-IRIS, (B) NTA, (C) MRPS, and (D) NFCM. Size distributions for U937 SEC EVs and 100K EVs (n=3 per group; with SD) for (E) ESP-IRIS, (F) NTA, (G) MRPS, and (H) NFCM. Please see Supplementary Figure 7 for graphs without error bars.

### Biological particle counting

Particle counts were next assessed. As before, we present the SP-IRIS data separately because this platform does not provide an overall particle count, but rather a number of events detected on surfaces printed with antibodies (shown here: to CD81 and to CD63 plus an isotype control). Consistent with expectations based on cellular tetraspanin expression and release, SP-IRIS showed that more H9 particles were captured by anti-CD81 than by anti-CD63 (Figure 5A) and that U937 particles could be captured by CD63 capture antibodies and not by CD81 capture antibodies (Figure 5B). For the remaining three platforms, which measure overall concentration, several trends were apparent (Figure 5C,D). First, for both the H9 and the U937 source, and for both EV separation methods, data were consistent with the results of SS counting in that NTA, NFCM, and MRPS measurements ordinally ranked from least particles/mL to greatest particles/mL. Second, MRPS and NFCM measured greater particle concentrations for 100K EVs than for SEC EVs (corrected for processing and dilution), although NTA results were similar. Finally, this is in contrast to results for the PS particles, where the three techniques returned roughly equivalent particle counts.

**Figure 5:**
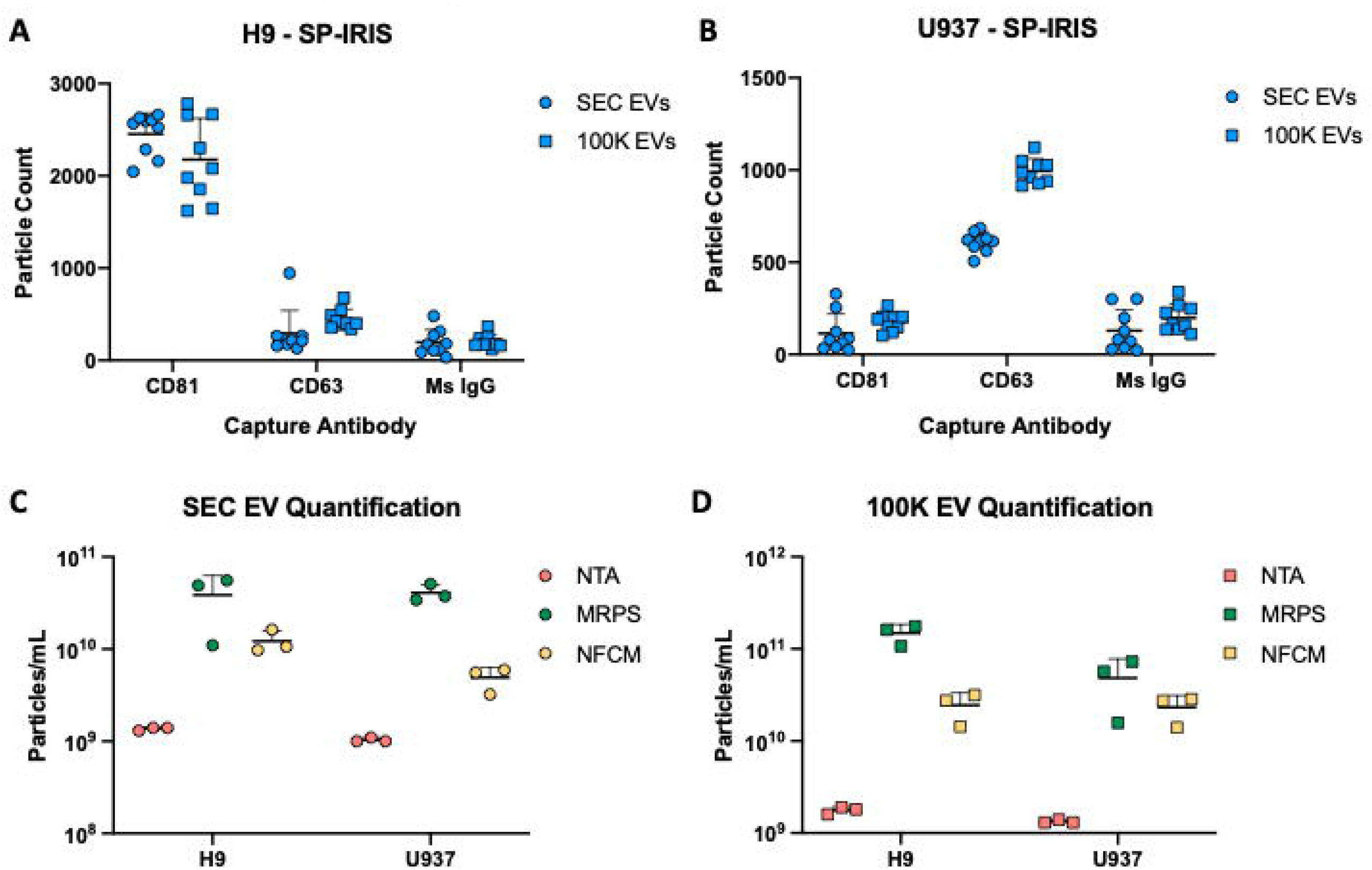
H9 and U937 particle quantification. SP-IRIS label-free capture for (A) H9 SEC EVs and 100K EVs and (B) U937 SEC EVs and 100K EVs using CD81, CD63, and mouse isotype control capture antibodies (measured on n=3 SP-IRIS chips and with n=3 antibody spots each; mean particle count/spot with SD). H9 and U937 particle quantification (n=3; mean particles/mL with SD) for (C) SEC EVs and (D) 100K EVs using NTA, MRPS, and NFCM.

### Do low concentrations of detergent affect MRPS measurements?

During the review process for this manuscript, a question arose about the possible effect of detergent on the reported results for biological particles measured by MRPS since a manufacturer-recommended dilution buffer contains 1% Tween 20. Specifically, it was proposed that the higher particle counts obtained for the same samples by MRPS and NTA could be due to artifactual small particle production when EVs are disrupted by detergent (see the comments section of [31]). We thus studied the effects of different concentrations of Tween 20 on MRPS measurements using archived aliquots of H9 EVs and U937 EVs. Because the maximum Tween 20 concentration used in any reported experiment was 0.5%, we conducted dilution series such that the same biological samples were measured in the presence of Tween 20 concentrations ranging from 0.1% to 0.9%. For reference, our highest concentration of Tween 20 was well below the concentrations previously reported to affect any of three classes of EVs [32]. Across Tween 20 concentrations, measured EV concentrations averaged 1.7×10^11^ +/-2.2×10^10^ (H9 EVs) and 1.5×10^11^ +/-2.9×10^10^ (U937 EVs), with no correlation between counts and detergent concentration (Supplementary Figure 8). Despite some variability in size profiles, there was also no evidence of a clear decrease in particle size with increasing detergent concentration (Supplementary Figure 8B,C).

### Single particle phenotyping by fluorescence

Fluorescence measurements for biological particles were done with three platforms. The MRPS platform has no fluorescence capabilities, so it was used only for sizing and counting. SP-IRIS performs a kind of single-particle phenotyping even in label-free mode, since diameter is measured for individual particles captured by antibodies and thus putatively positive for an antigen. What is more, captured particles can additionally be probed with fluorescently labeled antibodies. For chips incubated with H9 EVs (Figure 6A,B), EVs captured by CD81 antibodies were also generally positive for CD81 by fluorescence, and some also appeared to be CD63 positive. In contrast, CD63 capture spots were largely devoid of fluorescence for H9 EVs, as were (most) control capture spots. For chips incubated with U937 EVs (Figure 6C,D), events on CD63 capture spots were also positive for CD63 by fluorescence. CD81-linked fluorescence was at background levels for all spots. Note that numbers of “positive” events are higher in fluorescence mode than with label-free imaging (Figure 5A,B), likely, as discussed later, because fluorescence detection is more sensitive than reflectance imaging.

**Figure 6:**
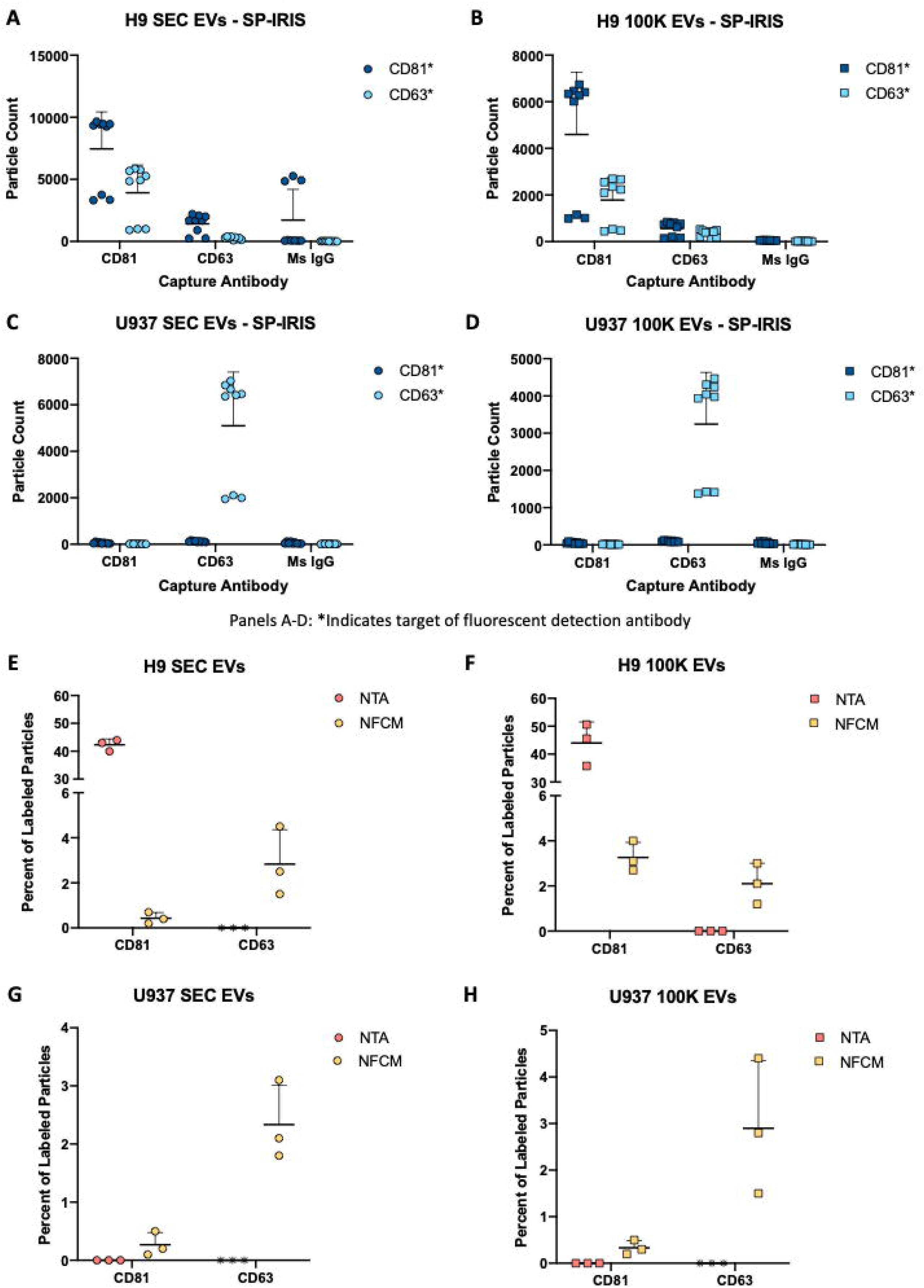
Particle phenotyping. SP-IRIS fluorescence detection using labeled anti-CD81 and anti-CD63 after particle capture with CD81, CD63, and mouse isotype control (n=3 per group; mean and SD) for (A) H9 SEC EVs, (B) H9 100K EVs, (C) U937 SEC EVs, and (D) U937 100K EVs. Percent of particles detected with fluorescently-labeled anti-CD81 and anti-CD63 by NTA and NFCM (n=3 per group; mean and SD) for (E) H9 SEC EVs, (F) H9 100K EVs, (G) U937 SEC EVs, and (H) U937 100K EVs. Asterisk. An asterisk indicates that, in the authors’ view, an antibody did not perform on the instrument; it does not necessarily mean that the antibody would not perform in another context or with additional optimization.

For the two remaining platforms with fluorescence capabilities, NTA and NFCM, results are shown as percent of total particles (Figure 6E-H). Approximately 40% to 50% of detected particles from H9 cells were positive for CD81 according to fluorescent NTA, while little to no CD81 signal was detected for U937 materials, consistent with expectations. However, we could not detect CD63-linked signal by fluorescent NTA for any sample. In contrast, NFCM detected either CD81 or CD63 on a small percentage of particles. The percentages were similar for the two tetraspanins for H9-derived particles. For U937 material, CD63-positive particles were more abundant than CD81-positive particles. No major differences between the SEC and 100K separation methods were apparent according to these data (Figure 6E-H).

## DISCUSSION

This study evaluated the abilities of four orthogonal technology platforms to size, count, and/or phenotype biological EVs and synthetic nanoparticles. Three of the technologies—SP-IRIS, NTA, and NFCM—are optical in nature and can perform some form of phenotyping/fluorescence analysis, while the other, MRPS, is an electric sensing platform that we did not attempt to apply to particle phenotyping. Although numerous comparisons of EV characterization platforms have been published previously [16,33–36], this study includes NFCM and MRPS and focuses in part on single-particle phenotyping.

### Detected particles: size-range sensitivity and refractive index matter

Whereas NTA, MRPS, and NFCM accurately and consistently measured the concentration of a known mixture of polystyrene particles, estimates of the number of silica particles varied substantially. NTA measured approximately 10-fold fewer SS particles than MRPS, while NFCM measured ∼ 3-fold fewer SS particles than MRPS. Since SS have a lower refractive index (n_SS_ ∼ 1.42 [37]) than PS (n_PS_ ∼ 1.59 [38]), one might predict that a mixture of EVs, with an even lower refractive index than silica [33,39], would have even larger variability between methods. Indeed, for EV preparations, average counts by NTA and MRPS differed by between one and two orders of magnitude. These outcomes emphasize that each platform has an effective range of measurement. MRPS is not sensitive to refractive index, but cartridges may clog (although we did not see evidence of this). In contrast, optical methods are quite sensitive to refractive index, and r^6 variation of scattered intensity limits dynamic range for a single instrument setting.

Thus, differences in output in part reflect different or overlapping particle populations that can be detected by the specific technologies, as indeed reported previously for several of these technologies [40]. That is, NTA and MRPS are similarly capable to detect a wide range of PS particle sizes. However, NTA may detect a more limited range of biological particles [41] than the MRPS platform using a small pore-size cartridge, in that MRPS may detect more of the smaller EVs along the power-law distribution. Signal for NTA and NFCM scales with radius to the 6^th^ power, which is general for light scattering in the Rayleigh approximation, whereas signal scales for MRPS and SP-IRIS with radius to the 3^rd^ power. Thus, because of finite dynamic range, NTA will be biased to detecting fewer of the small particles in a sample compared with MRPS. Of course, this will also depend on how the NTA the instrument and analysis settings are configured. One might over-expose the large particles in order to see the small ones, for example, or to increase sensitivity and maximize counts, but the outcome of this adjustment may be limited by glare from the large particles.

### Is it important to resolve different particle size populations?

SP-IRIS, MRPS, and NFCM could resolve up to four populations of synthetic nanoparticles with different diameters. We note that distinct populations were somewhat obscured when MRPS results were averaged for SS, but not for PS – see also Supplementary Figure 6 – which may reflect aggregation of the SS due to the electrolyte solution (PBS) required for MRPS and the convolution of experimental uncertainties in particle concentration and size measurements. Alternatively, as suggested by an astute reviewer, a PS standard could be run later on the same sample for scaling of size and concentration. Also noteworthy is that the NFCM platform distinguished subpopulations of SS particles quite well, but that this is likely because the same beads are used to calibrate the instrument. While detecting the expected concentration of high refractive index PS particles, NTA was unable to resolve individual particle populations and instead characterized the SS particles as a broad population distribution centered on an “average” size. To be sure, it may be possible to resolve discretely sized particle populations using NTA with mixtures at different ratios of sizes. We could not do so with the mixtures we used. Whether this matters for biological particles is unclear. It does not seem that biological samples would contain unique EV subpopulations with exquisitely defined sizes, except perhaps for samples from sources infected with specific enveloped viruses. NTA does seem to be capable of detecting shifts in population distributions, and this capability might be more important for biological particles than resolving subpopulations.

### On counting by SP-IRIS/fluorescence

In this study, neither SP-IRIS label-free measurements nor subsequent fluorescence detection could be used directly to estimate overall particle concentration. Instead, SP-IRIS is best used to understand ratios within populations and for single-particle phenotyping. Only a subset of EVs bind to any given affinity reagent “spot.” Binding is determined by diffusion (which is slow for EVs), presence and density of recognized surface markers, and affinity characteristics of antibody-to-antigen binding. The bound population of particles remaining after wash steps is only a small proportion of the total in the input material and cannot be used to determine overall concentration. Interestingly, fluorescence results often indicated higher particle concentrations than returned by label-free counting, even though particles positive for a particular antigen are expected to be only a subset of the captured population (different antigens) or to approach equality (if the capture antigen is targeted and antigen is abundant). Counts are higher because fluorescence detection is more sensitive than label-free. That is, fluorescence detects positive particles that may be below the limit of label-free detection. Also of note, capturing EVs onto the chip via surface markers may render those markers less available to subsequent binding by fluorescently-labeled antibodies. For example, in Figure 6A and B, CD81-captured particles that also display CD63 appear to have more CD63 available for binding by fluorescent antibodies compared with CD63+ particles that have had at least some portion of their CD63 sequestered by the surface-bound antibodies.

### Did any platforms identify differences between EV separation technologies?

For both biological sources of EVs, we used two methods of EV separation: dUC (100K EVs), which has been the most common method for EV separation [42–44], and a combination of filtration and size exclusion chromatography (SEC EVs) [45]. According to some evidence in the literature, dUC leads to more protein contamination and aggregation and damage of EVs [46– 48]. It should be noted that alternative viewpoints can also be found [18]. However, protein particle contamination might be expected to introduce more and smaller particles. This outcome is indeed observed based upon TEM background and particle profile shifts towards smaller particles for several of the platforms. On the other hand, evidence of aggregation by dUC is not apparent in the data presented here. We cannot rule out aggregation, however, only that the techniques used here did not appear to detect it; we also do not wish to put too fine a point on these comparisons, which are based on limited data.

### Single-particle phenotyping

For the three techniques with single-particle phenotyping capabilities (SP-IRIS, NTA, and NFCM), each has advantages and challenges. SP-IRIS was able to achieve true “multiplexed” detection, in that signal could be obtained above background for up to three fluorescent channels. At the time of our evaluations, the NTA platform we used could not perform simultaneous multi-channel measurements and thus was not a true single-particle multiplexing platform. Instead, sequential filter switches were required, such that the same particles could not be tracked in different channels. Finally, although the nano-flow technology may be capable of multiplexed phenotyping, we did not explore this capability here.

### Summary of findings: Table 2

In Table 2, we attempt to summarize our findings and views about the four investigated techniques. **Detectable size ranges** for biological particles: these should be considered to be rough estimates. If we accept the assumption that EVs follow a power-law size distribution (the smaller, the more abundant, with lower bounds defined by membrane curvature constraints), then no evaluated platform effectively detects the very smallest particles. However, SP-IRIS, MRPS, and NFCM appear to detect slightly smaller particles than NTA **under the conditions and settings we tested**. For NTA, MRPS, and NFCM, linear ranges for **particle concentration** for all instruments begin around 1×10^7^ particles/mL (or slightly lower) and extend from about one order of magnitude (NTA) to multiple orders of magnitude (MRPS). This spread is important, since the wider the range, the fewer time-consuming concentrations or dilutions must be done to place an unknown particle population into the measurable range. SP-IRIS is a special case, since particles are captured by affinity, and overall concentration cannot easily be estimated. In our hands, particle concentrations must be high (>>1×10^7^ particles /mL) even for abundant antigens. Furthermore, the optimal captured particle counts are roughly several thousand per antibody spot (although this may vary). To hit a tight “sweet spot”, trial dilutions may be needed. Furthermore, the optimal dilution may differ for different antibodies on the chip because of different percentages of EVs positive for a particular antigen, per-EV antigen abundance, and antibody performance. Hence, dilutions are usually most important and potentially time-consuming for SP-IRIS. Related to dilution is the **volume of input material** required for a single reading. Assuming each platform is provided with a suspension at 1×10^7^ particles per mL, the required volume of a dilution at this concentration ranges from 5 µL (MRPS) to around 1 mL (NTA). Of course, the actual volume/number of EVs needed will also depend on the number of concentrations/dilutions required to reach the measurable concentration range. The input volume difference is also inconsequential for highly abundant materials, but may be important for low-abundance EV samples. **If done, optional calibration steps** are rapid for NTA and MRPS (around 20 minutes). For NFCM, we find that calibration can be as short as 20 minutes but can sometimes take longer. Time for sample dilutions is most difficult to estimate, but is expected to correlate inversely with the range of measurement for each platform. **Read time** ranges from five minute to about half an hour per sample. Note that the times we indicate are for sizing and counting only. Optional fluorescence measurements for the relevant platforms would in some cases add processing time for antibody incubations and removal, as well as for read times (except for NFCM). For SP-IRIS, we should also note that, although the total hands-on and read time is longer than for other techniques, each reading includes on-chip replicates, multiple capture antibodies, and up to three fluorescence readouts per capture antibody.

**Table 2.**
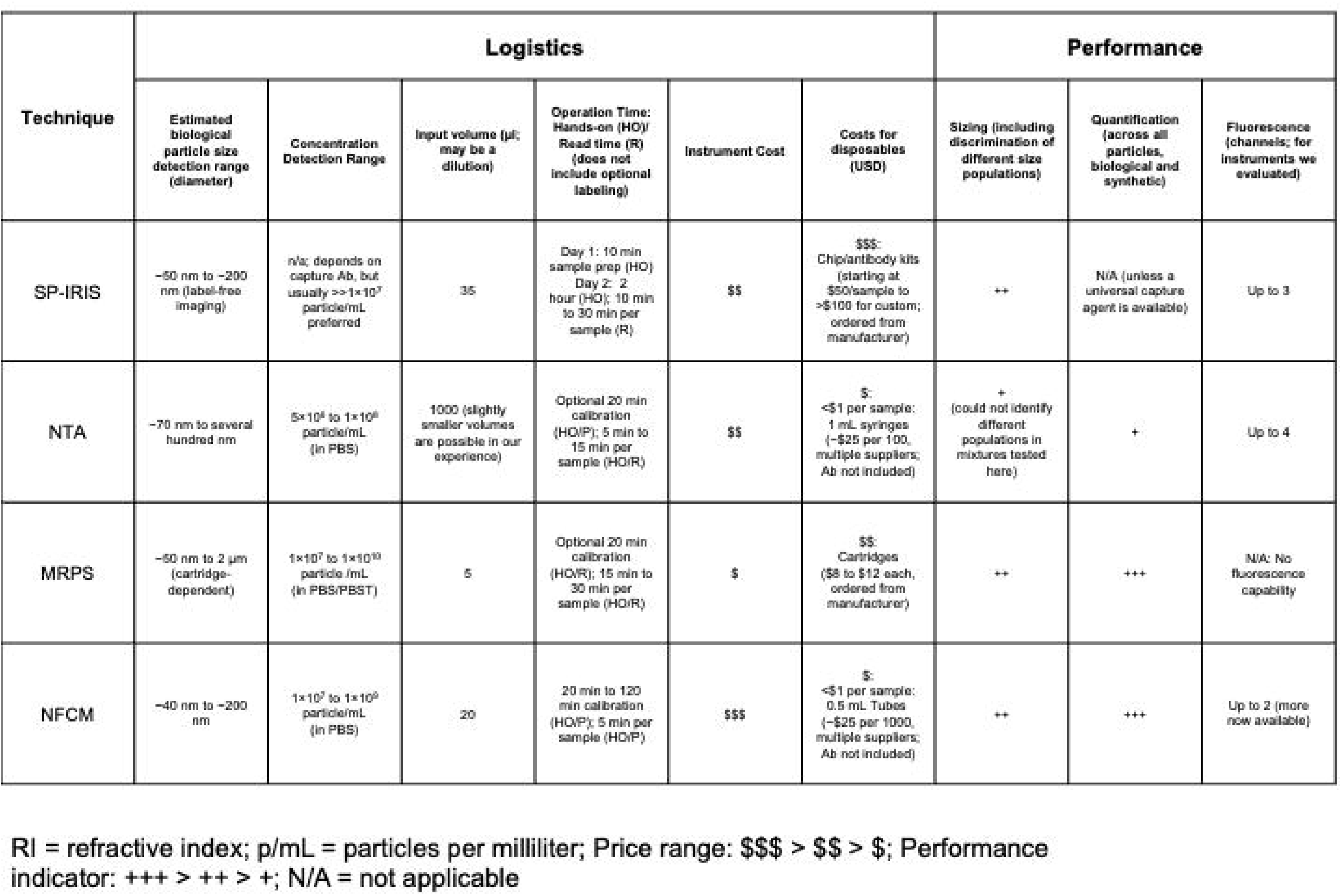
Evaluation summary

***Costs*** for the platforms include initial outlay, disposable costs, and maintenance costs. For acquisition, the MRPS system is most economical, while the NFCM platform is the most expensive. For basic counting and sizing, operating costs for NTA and NFCM are negligible. Adding optional fluorescence increases these costs by amounts that are antibody-dependent. The MRPS system uses disposable cartridges that currently cost USD 8 to USD 12 each. The SP-IRIS platform has the highest disposable costs, with each sample requiring at least one chip that costs from USD 50 to >USD 100 each. Since optimal dilutions must be made and may be different for different capture materials on the same chip, multiple chips may be needed for the same sample. Chips also cannot be chemically stripped and re-used, at least not in our hands (Mallick and Witwer, unpublished data). Shelf-life of the chips is also a consideration. However, the company’s development of chips with extended shelf-life may overcome this potential hindrance. As noted, though, under optimal conditions, the platform provides multi-dimensional information that may justify costs and logistical challenges. We should also mention that chips for the SP-IRIS and MRPS instruments are currently available only from the instrument manufacturer for that particular measurement technique. As for maintenance costs, we are unable to estimate them at this time for any platform.

There are several limitations of our study as well as questions that might be investigated in the future. Since several of the platforms easily distinguished multiple populations of synthetic particles, but did not always identify the expected size for each population, normalization strategies including spiked-in standards could be useful. However, a question we have not addressed here is how components of the suspension medium might affect spiked-in synthetic material. For example, if synthetic beads are spiked into biological fluids, will they acquire “coronas” that change their measured properties? Our study, by examining only several distinct sizes of synthetic particles, also does not rigorously define the range of size detection for biological particles for each platform. Likewise, our estimates of range of concentration measurement for each instrument are simplistic. In theory, a “single particle” detection instrument is capable of detecting a single particle, although measurement noise, contaminants, and the time required to “find” the single particle are real-world considerations that make this unlikely.

## In conclusion

- For any platform and configuration, particle counting is accurate only within a certain range. Sensitivity for particles of different sizes and refractive indices should be considered. Recall that signal for light scattering methods like NTA and NFCM scales with radius to the sixth power, while signal for SP-IRIS and MRPS scales with radius to the third power. However, in our hands, the NFCM platform is more sensitive than NTA for small and low refractive index particles.
- Different size populations within a mixture of synthetic nanoparticles can be identified by SP-IRIS, MRPS, and NFCM, but not, in our hands, by NTA. The individual sizes are not always accurately assigned, however, emphasizing the importance of calibration.
- SP-IRIS, NTA, and NFCM offer fluorescent particle phenotyping, while MRPS does not. Multiplexed biological particle phenotyping of tetraspanins was easily achieved with the SP-IRIS platform (one-antibody capture and up to three-antibody fluorescence detection).
- Appropriate reference materials are needed for better evaluation of single particle phenotyping capabilities, including multiplexed phenotyping.
- Rather than relying on a single platform, consider using orthogonal technologies.
- Both acquisition and recurring costs should be considered before choosing a platform.
- No evaluated platform is necessarily “better” or “worse” than others; rather, it is important to be aware of the capabilities of each platform with respect to each particle population of interest and the population attributes that are of greatest interest.

## Acknowledgements

The authors thank members of the Witwer Laboratory and the Retrovirus Laboratory and various members of the International Society for Extracellular Vesicles for discussions and support. The authors acknowledge NanoView Biosciences, Particle Metrix, Spectradyne, and NanoFCM representatives for discussions, for arranging demonstrations, and for helpful comments on the preprint. Electron microscopy images were acquired in the Johns Hopkins University School of Medicine Institute for Basic Biomedical Sciences Microscope Facility.

---

**Supplementary Figure 1:**
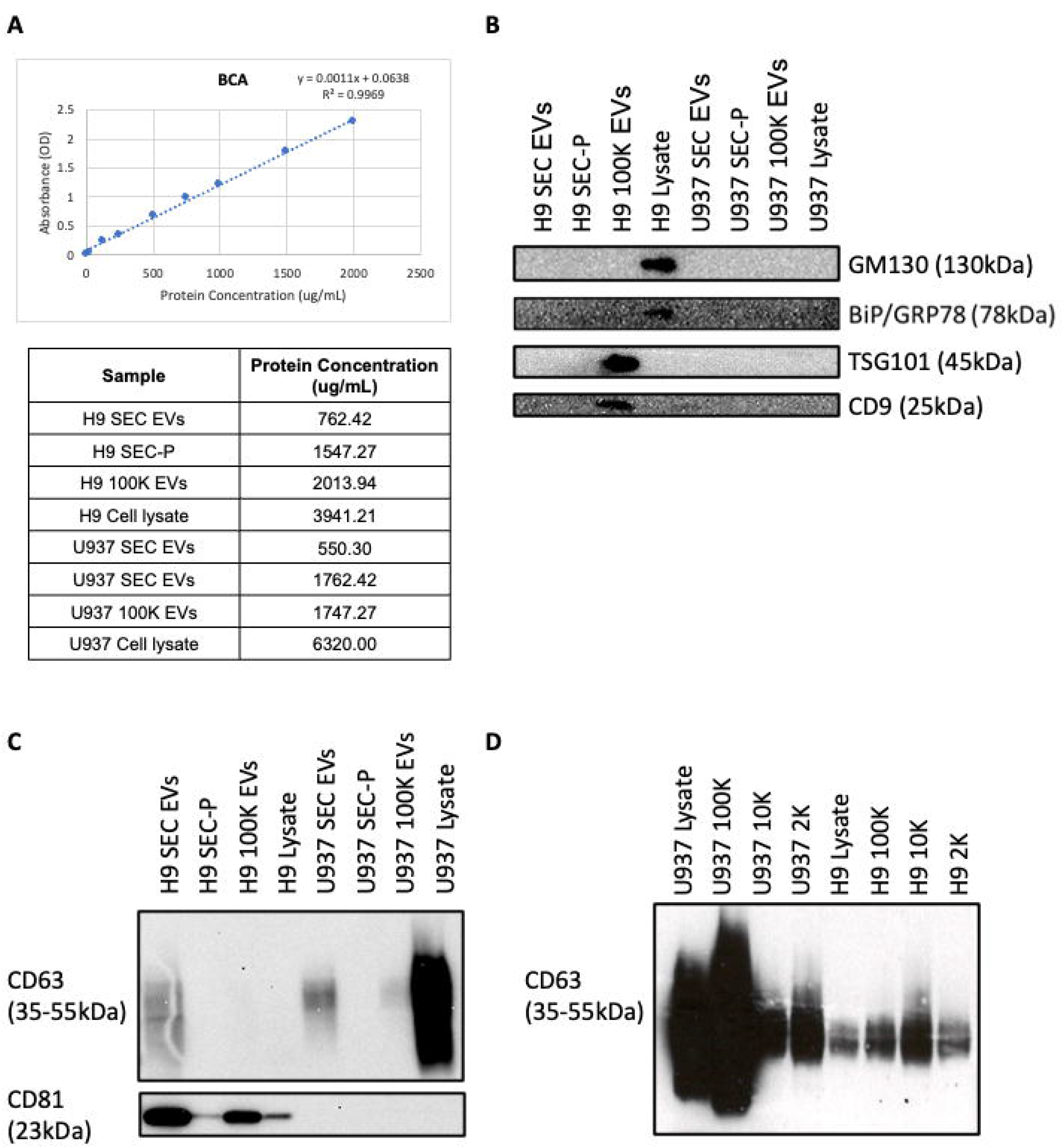
Additional EV characterization. (A) Representative BCA assay (protein concentration) results from one batch of EV separations. (B) Immunoblot analysis of separated EVs using the same samples as in Figure 1 but probing for GM130 and BiP/GRP78 (expected to be depleted in EVs) and TSG101 and CD9 (expected to be enriched in EVs). (C) An immunoblot from a previous experiment that was shown in Figure 1 of a previous version of this manuscript. (D) Overexposed CD63 results from a previous set of EV separations from U937 and H9 cells (here, using differential ultracentrifugation at 2K × g, 10K × g, and 100K × g) showing that, in some experiments, CD63 is indeed enriched in the 100K EV-enriched pellet, and that lengthy exposure confirms the presence of CD63 in H9 cell lysate, albeit at low levels.

**Supplementary Figure 2:**
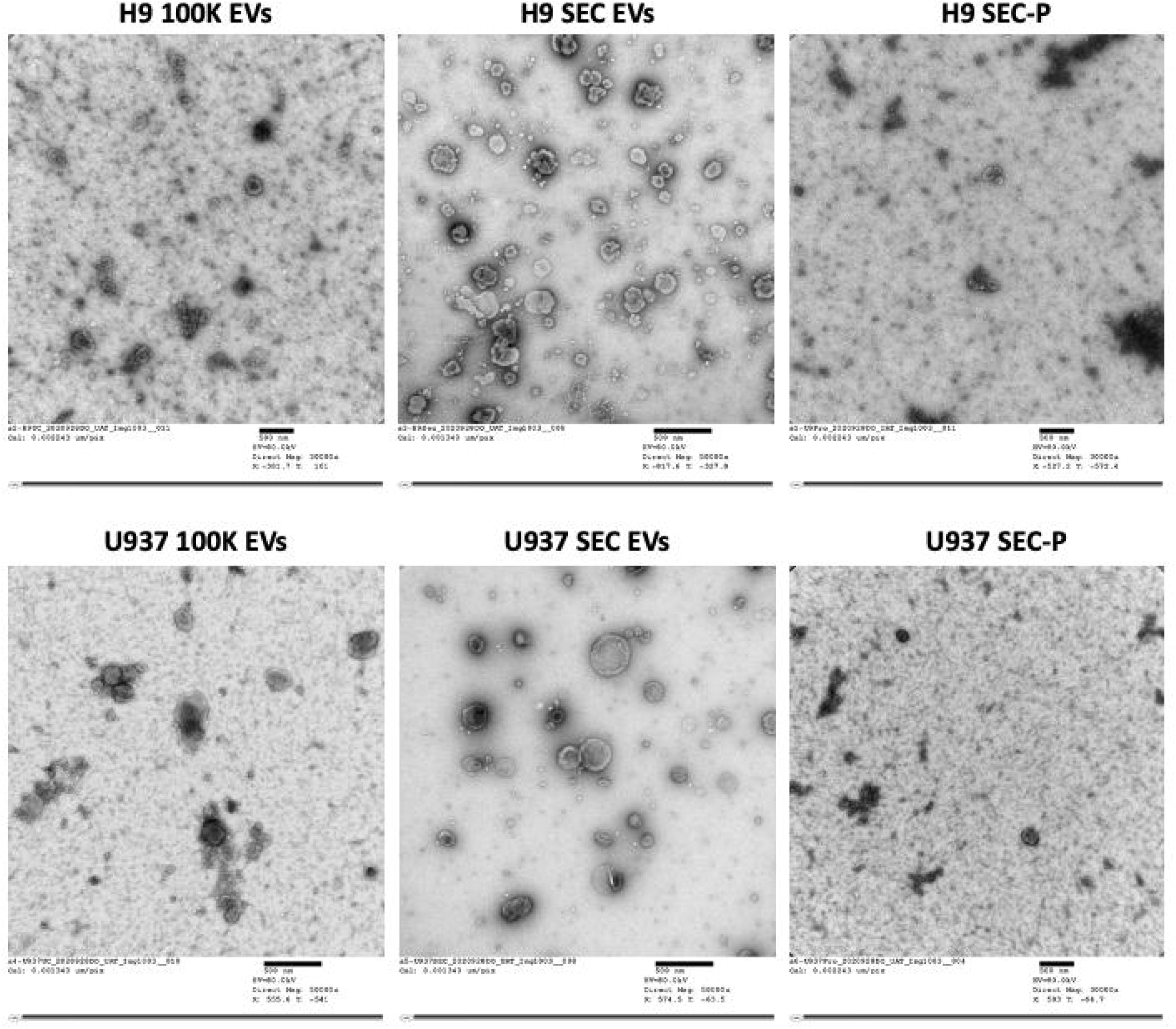
Additional TEM of EVs and SEC protein fractions. Scale bars, as indicated, are 500 nm.

**Supplementary Figure 3:**
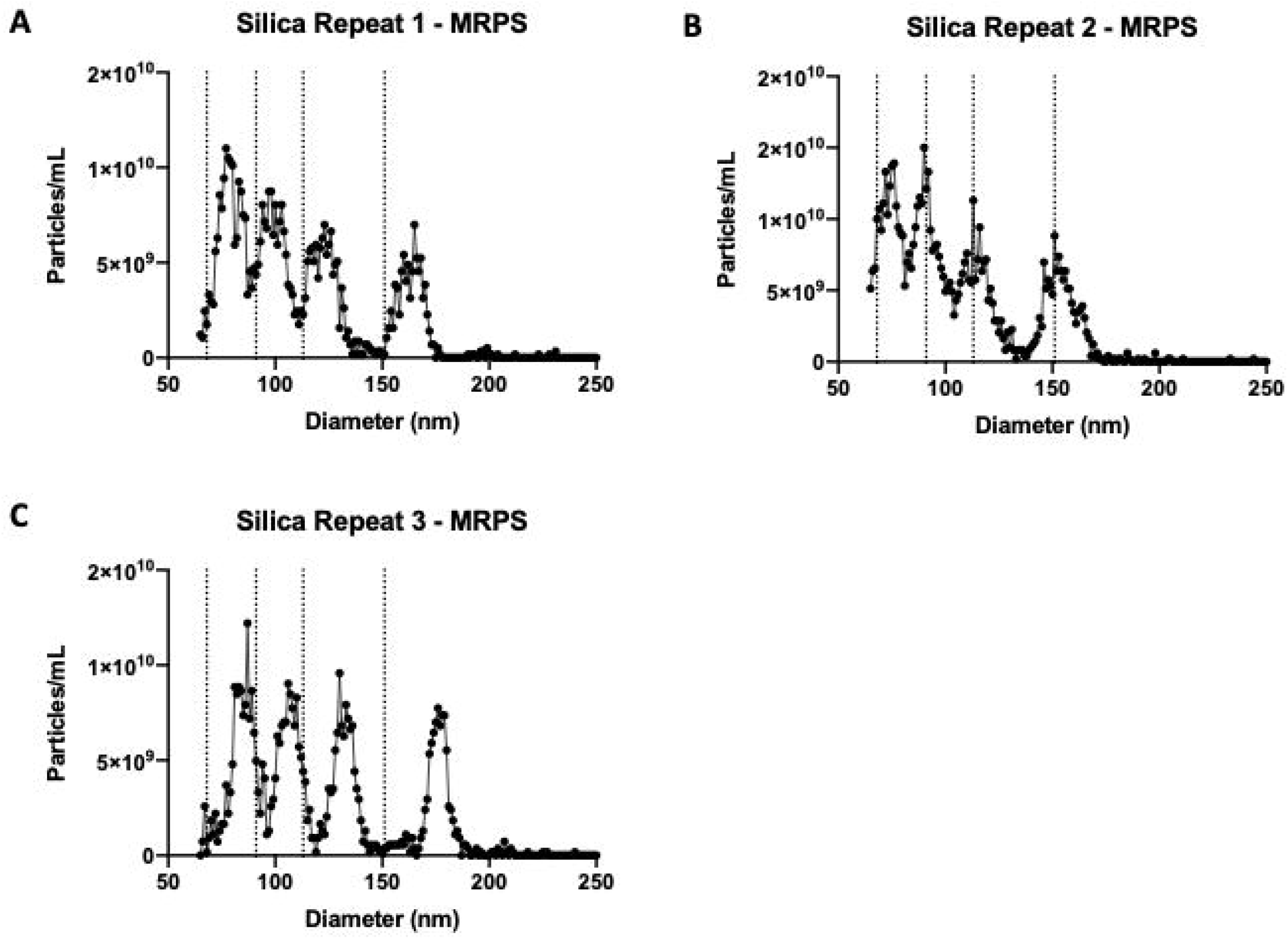
Individual SS measurements by MRPS. Repeat 1 (A) can also be found as an inset in Figure 2C. (B) and (C) are additional repeats using the same SS mixture.

**Supplementary Figure 4:**
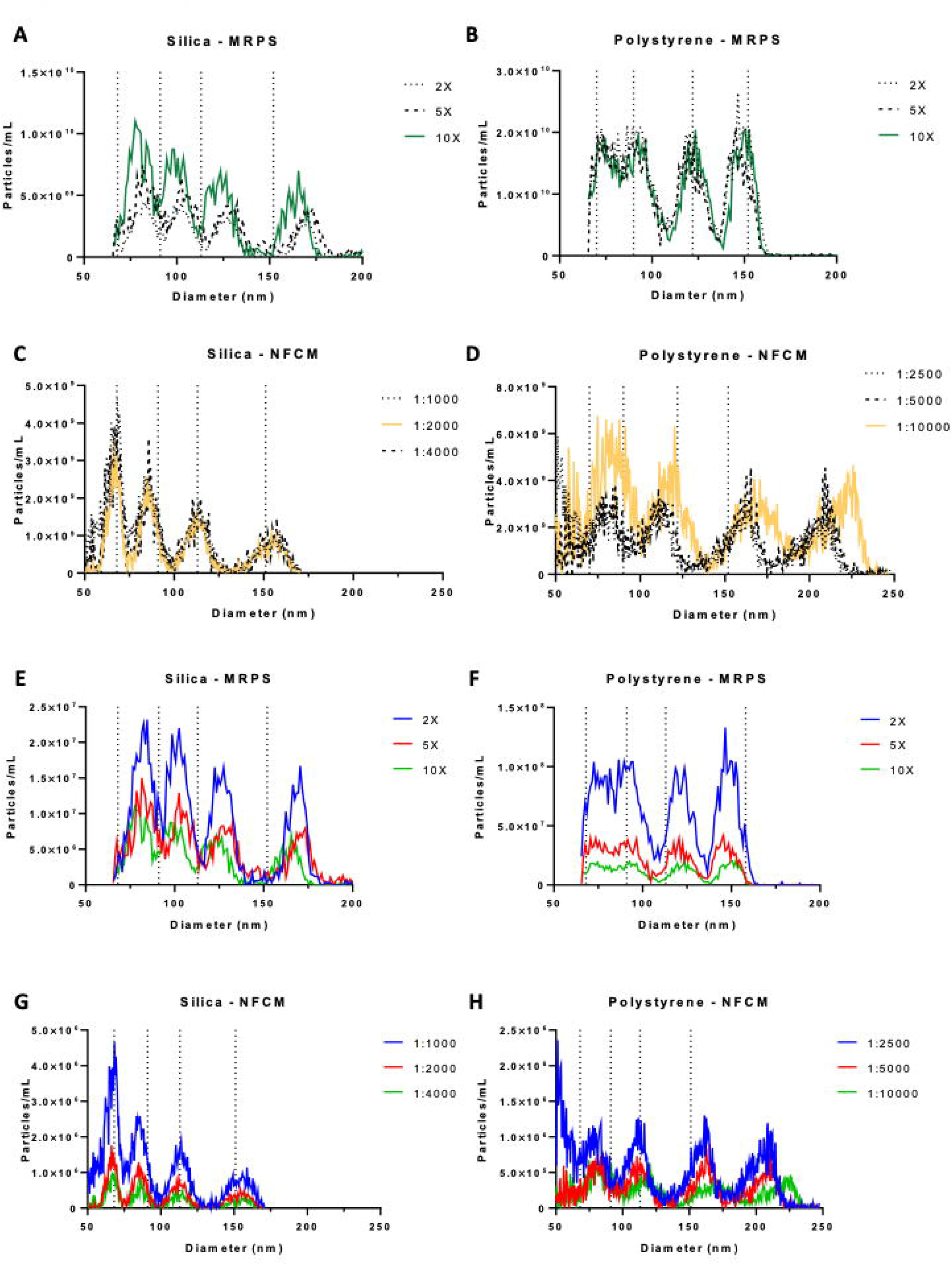
MRPS and NFCM dilution series. SS (A) and PS (B) were diluted 2×, 5×, and 10× by volume to determine the optimal dilution for NTA analysis. SS (C) were diluted 1:1000 (v:v), 1:2000 (v:v), and 1:4000 (v:v) and PS (D) were diluted 1:2500 (v:v), 1:5000 (v:v), and 1:10000 (v:v) to determine the optimal dilution for NFCM analysis. The upper four panels are dilution-corrected data. Bottom four panels: raw data for SS (E) and PS (F) on MRPS and SS (G) and PS (H) on NFCM. Optimal dilutions are indicated by green or yellow (MRPS and NFCM, respectively).

**Supplementary Figure 5:**
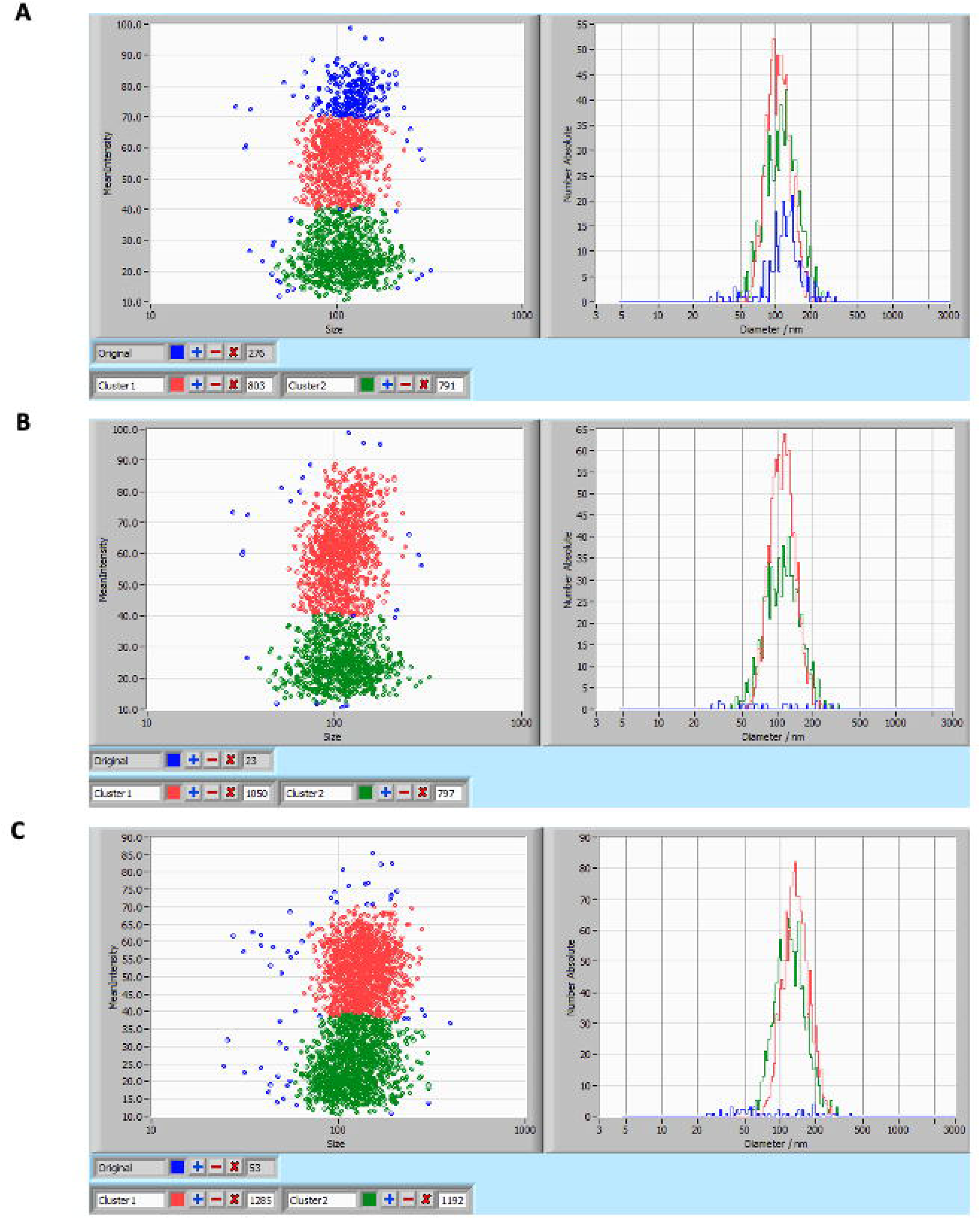
Intensity gates for assessment of populations by NTA. Data from NTA measurements of PS (A, B) and SS (C) were used to assign gates based on intensity. Note that (A) and (B) are the same data with different gates. Left panels are intensity vs diameter plots. Right panels are abundance vs diameter for each indicated (color-coded) intensity gate.

**Supplementary Figure 6:**
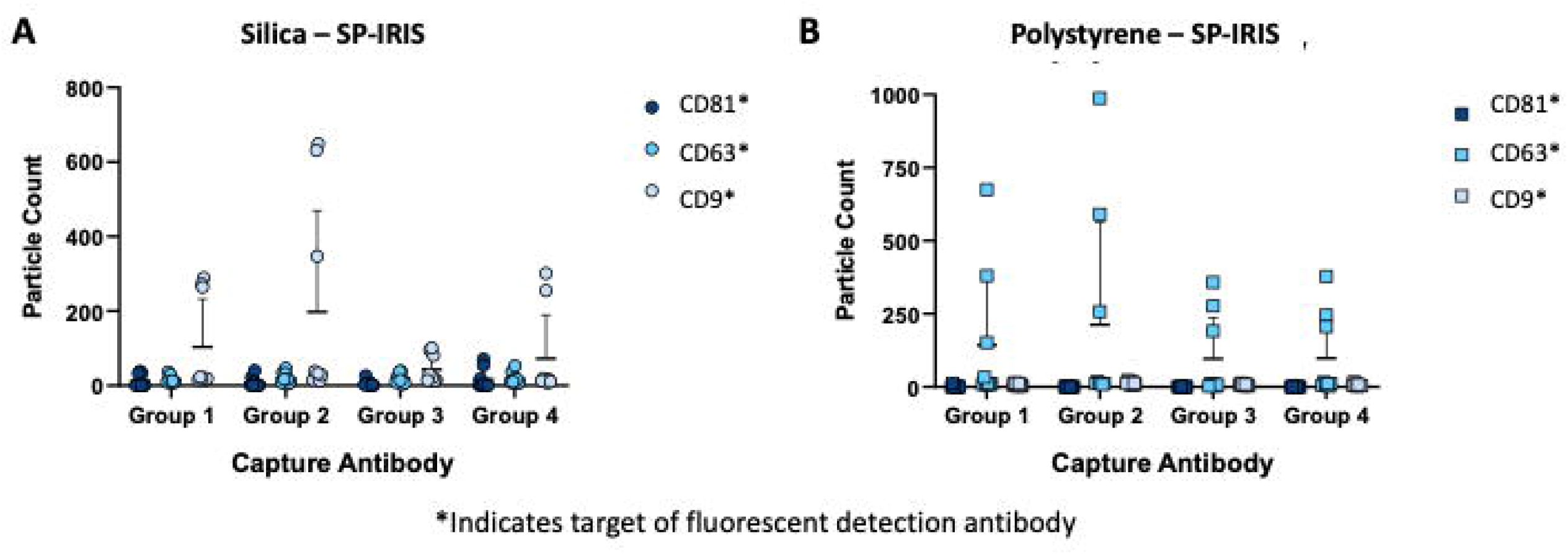
SP-IRIS background fluorescence for SS and PS. SP-IRIS fluorescence detection using fluorescently labeled anti-CD81, anti-CD63, and anti-CD9 after drying (A) SS and (B) PS onto SP-IRIS chips and measuring particles dried onto spots corresponding to the four antibody groups (n=3 chips per group and 3 spots per antibody per chip; mean and SD are indicated by bars and whiskers).

**Supplementary Figure 7:**
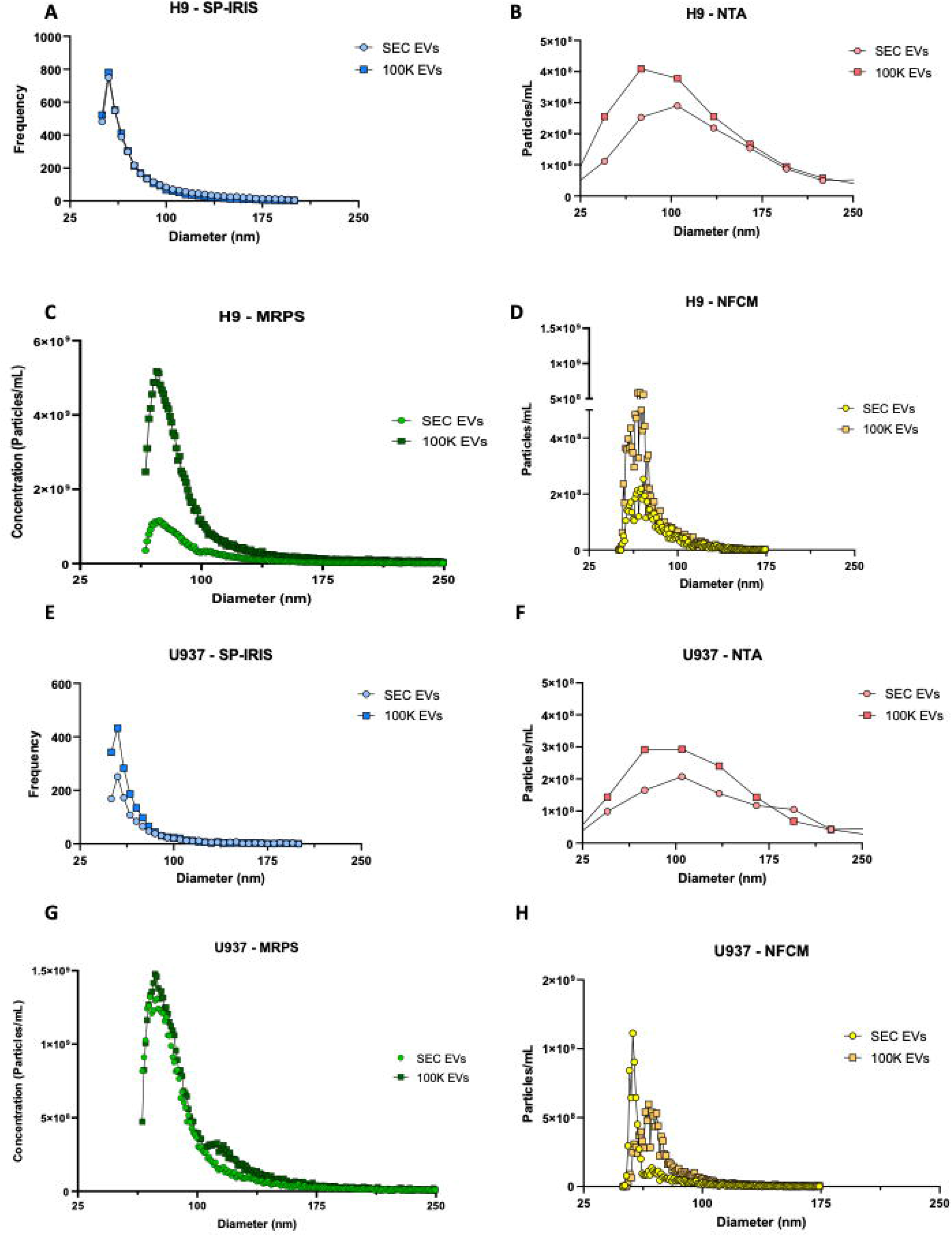
H9 and U937 EV size distribution, no error bars. This figure depicts the same data as shown in Figure 4, but without error bars for clarity.

**Supplementary Figure 8:**
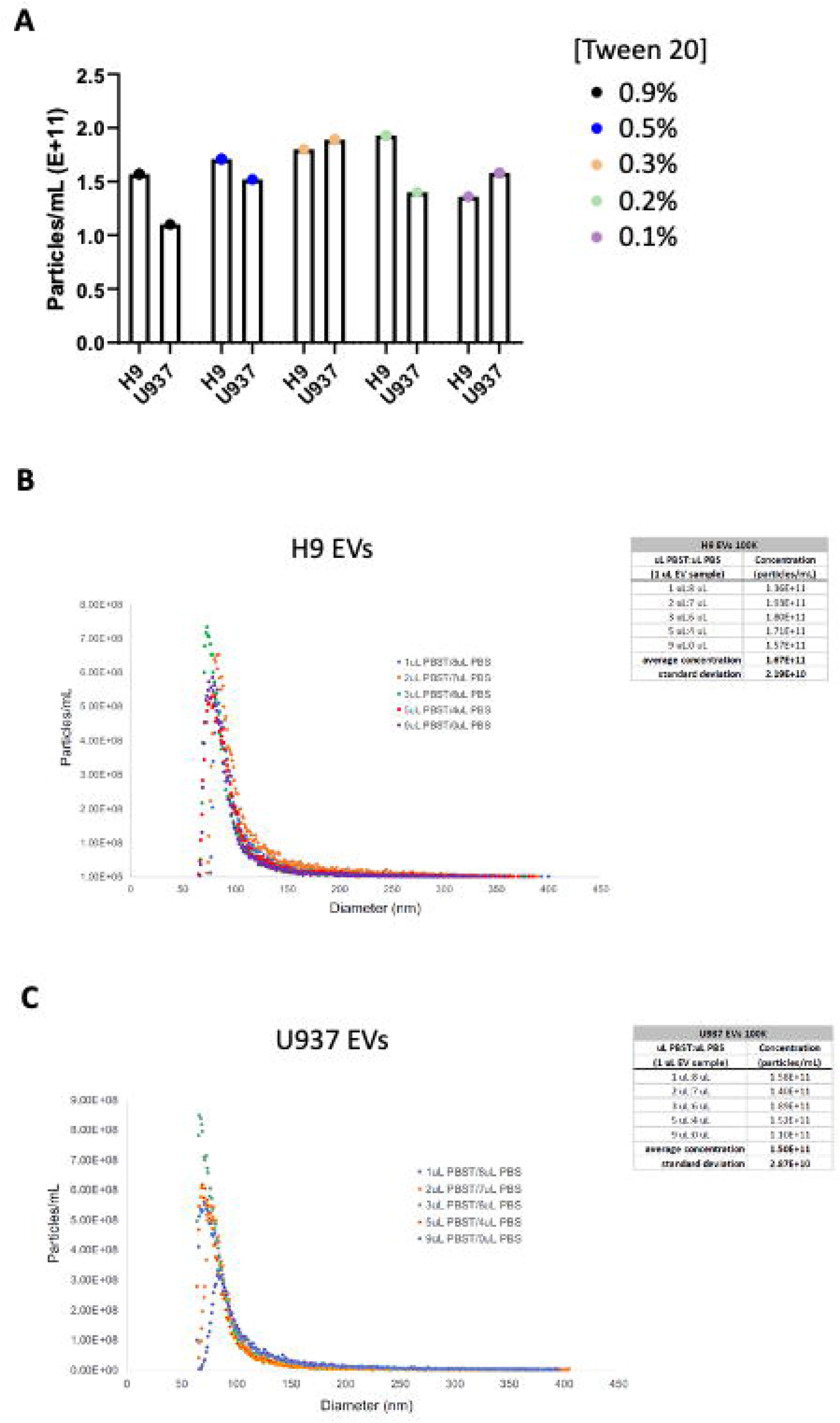
Effects of Tween 20 on MRPS measurements. H9 and U937-derived EVs were mixed with Tween 20 to final concentrations ranging from 0.1% to 0.9% and measured by MRPS. (A) Particle counts. (B) and (C) depict size distributions for H9 and U937 EVs, respectively, along with insets displaying concentrations.

**Supplementary Table 1.**
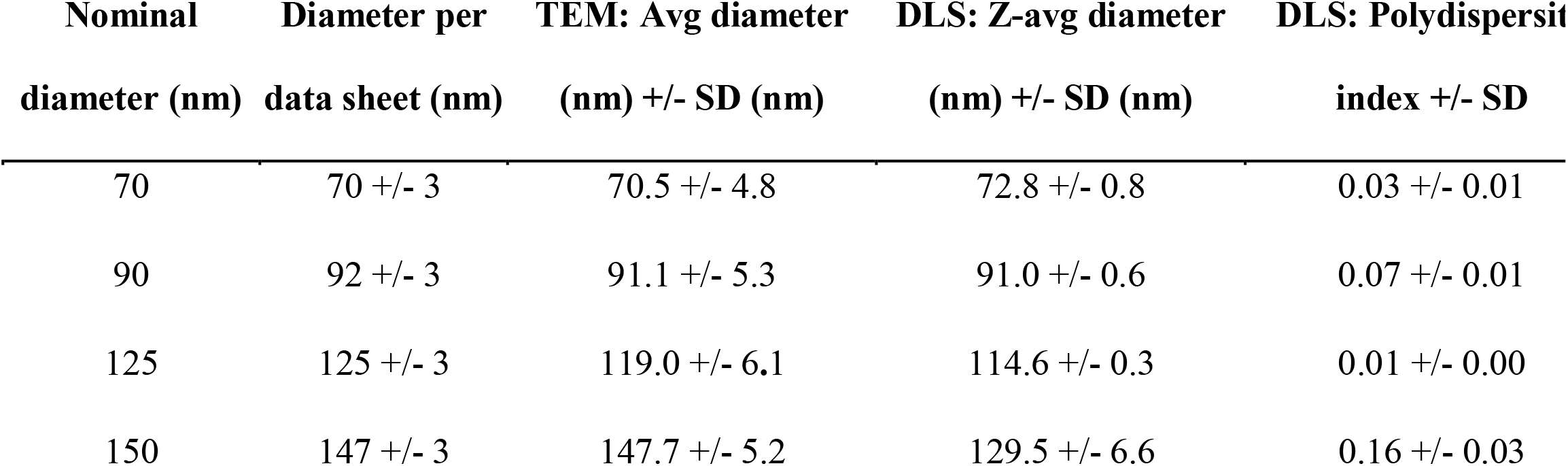
TEM and DLS measurements of selected PS beads. Three preparations of each PS bead population were measured three times each. Avg = arithmetic mean; SD = standard deviation

**Supplementary Table 2:**
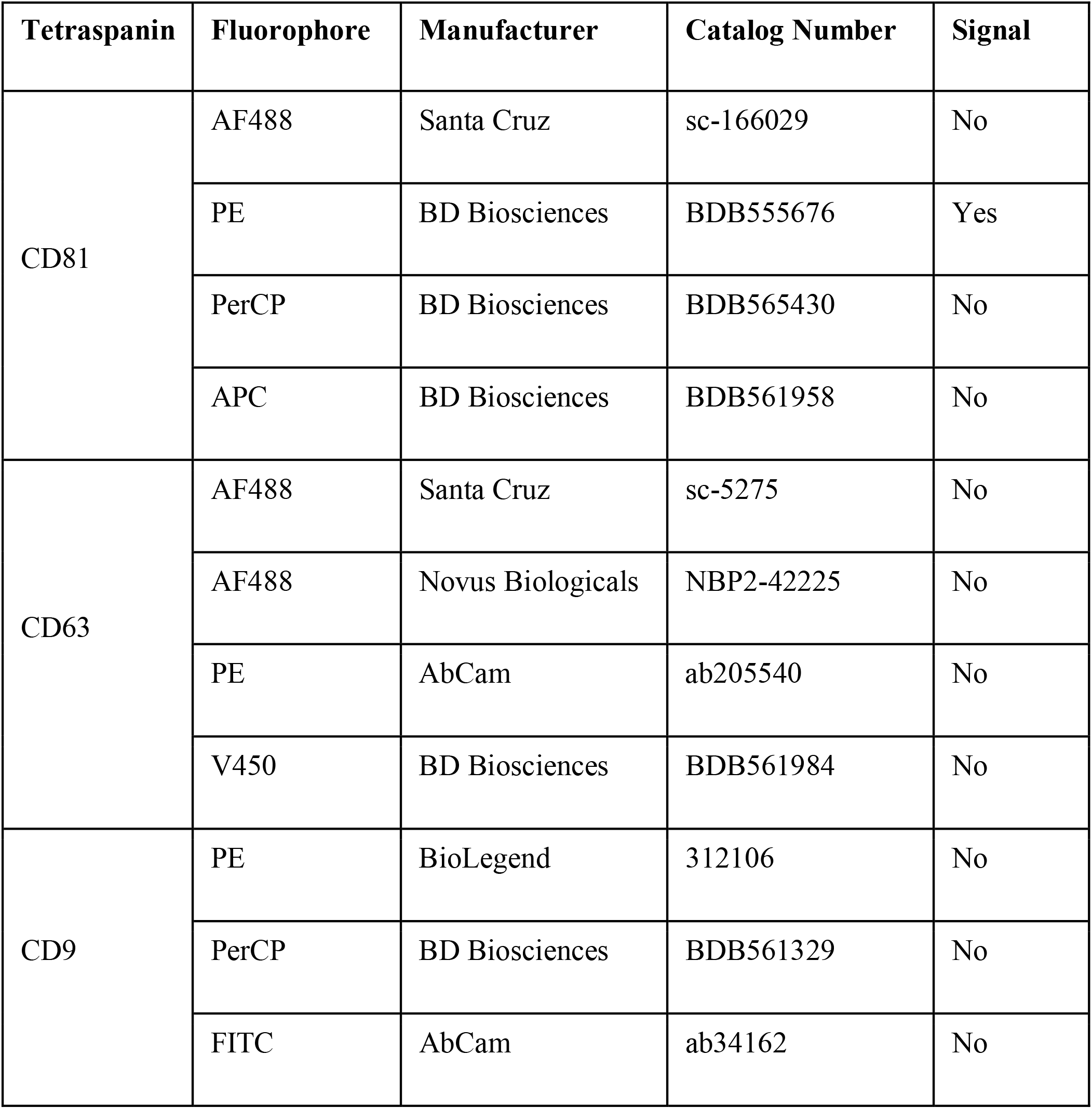
Antibodies tested with fluorescent NTA. H9 100K EVs were diluted 1:1 (v:v) in PBS. 9 µL of diluted EVs were mixed with 1 µL of antibody and incubated for 2 hours at room temperature. Samples were then diluted 1:1000 and measured in scatter and fluorescent modes using NTA. We would like to stress that our inability to obtain signal with these antibodies likely indicates that further optimization is needed, not necessarily that the antibodies are unsuited to this use.

| Tetraspanin | Fluorophore | Manufacturer | Catalog Number | Signal |
| --- | --- | --- | --- | --- |
| CD81 | AF488 | Santa Cruz | sc-166029 | No |
|  | PE | BD Biosciences | BDB555676 | Yes |
|  | PerCP | BD Biosciences | BDB565430 | No |
|  | APC | BD Biosciences | BDB561958 | No |
| CD63 | AF488 | Santa Cruz | sc-5275 | No |
|  | AF488 | Novus Biologicals | NBP2-42225 | No |
|  | PE | AbCam | ab205540 | No |
|  | V450 | BD Biosciences | BDB561984 | No |
| CD9 | PE | BioLegend | 312106 | No |
|  | PerCP | BD Biosciences | BDB561329 | No |
|  | FITC | AbCam | ab34162 | No |

## REFERENCES

[1] Théry C, Witwer KW, Aikawa E, et al. Minimal information for studies of extracellular vesicles 2018 (MISEV2018): a position statement of the International Society for Extracellular Vesicles and update of the MISEV2014 guidelines. J Extracell Vesicles 2018;7:1535750.

[2] Yáñez-Mó M, Siljander PR-M, Andreu Z, et al. Biological properties of extracellular vesicles and their physiological functions. J Extracell Vesicles 2015;4:27066.

[3] Witwer KW, Théry C. Extracellular vesicles or exosomes? On primacy, precision, and popularity influencing a choice of nomenclature. J Extracell Vesicles 2019;8:1648167.

[4] Cocucci E, Meldolesi J. Ectosomes and exosomes: shedding the confusion between extracellular vesicles. Trends Cell Biol 2015;25:364–72.

[5] Lázaro-Ibáñez E, Lässer C, Shelke GV, et al. DNA analysis of low- and high-density fractions defines heterogeneous subpopulations of small extracellular vesicles based on their DNA cargo and topology. https://DoiOrg/101080/2001307820191656993 2019.

[6] Russell AE, Sneider A, Witwer KW, et al. Biological membranes in EV biogenesis, stability, uptake, and cargo transfer: an ISEV position paper arising from the ISEV membranes and EVs workshop. J. Extracell. Vesicles, vol. 8, Taylor and Francis Ltd.; 2019.

[7] Tkach M, Kowal J, Théry C. Why the need and how to approach the functional diversity of extracellular vesicles. Philos Trans R Soc Lond B Biol Sci 2018;373:20160479.

[8] Lässer C, Jang SC, Lötvall J. Subpopulations of extracellular vesicles and their therapeutic potential. Mol Aspects Med 2018;60:1–14.

[9] Lanier LL, Engleman EG, Gatenby P, et al. Correlation of functional properties of human lymphoid cell subsets and surface marker phenotypes using multiparameter analysis and flow cytometry. Immunol Rev 1983;74:143–60.

[10] Reddington AP, Trueb JT, Freedman DS, et al. An Interferometric Reflectance Imaging Sensor for Point of Care Viral Diagnostics. IEEE Trans Biomed Eng 2013;60:3276–83.

[11] Lopez CA, Daaboul GG, Vedula RS, et al. Label-free multiplexed virus detection using spectral reflectance imaging. Biosens Bioelectron 2011;26:3432–7.

[12] Daaboul GG, Gagni P, Benussi L, et al. Digital Detection of Exosomes by Interferometric Imaging. Sci Rep 2016;6:37246.

[13] Dragovic RA, Gardiner C, Brooks AS, et al. Sizing and phenotyping of cellular vesicles using Nanoparticle Tracking Analysis. Nanomedicine 2011;7:780–8.

[14] Sokolova V, Ludwig AK, Hornung S, et al. Characterisation of exosomes derived from human cells by nanoparticle tracking analysis and scanning electron microscopy. Colloids Surf B Biointerfaces 2011;87:146–50.

[15] Giebel B, Helmbrecht C. Methods to analyze EVs. Methods Mol. Biol., vol. 1545, Humana Press Inc.; 2017, p. 1–20.

[16] Anderson W, Kozak D, Coleman VA, et al. A comparative study of submicron particle sizing platforms: Accuracy, precision and resolution analysis of polydisperse particle size distributions. J Colloid Interface Sci 2013;405:322–30.

[17] Fraikin JL, Teesalu T, McKenney CM, et al. A high-throughput label-free nanoparticle analyser. Nat Nanotechnol 2011;6:308–13.

[18] Tian Y, Gong M, Hu Y, et al. Quality and efficiency assessment of six extracellular vesicle isolation methods by nano-flow cytometry. J Extracell Vesicles 2020;9.

[19] Tian Y, Ma L, Gong M, et al. Protein Profiling and Sizing of Extracellular Vesicles from Colorectal Cancer Patients via Flow Cytometry. ACS Nano 2018;12:671–80.

[20] Ahn JY, Datta S, Bandeira E, et al. Release of extracellular vesicle miR-494-3p by ARPE- 19 cells with impaired mitochondria. Biochim Biophys Acta Gen Subj 2020:129598.

[21] Gámbaro F, Li Calzi M, Fagúndez P, et al. Stable tRNA halves can be sorted into extracellular vesicles and delivered to recipient cells in a concentration-dependent manner. RNA Biol 2019.

[22] Huang Y, Cheng L, Turchinovich A, et al. Influence of species and processing parameters on recovery and content of brain tissue-derived extracellular vesicles. J Extracell Vesicles 2020.

[23] Van Deun J, Mestdagh P, Agostinis P, et al. EV-TRACK: Transparent reporting and centralizing knowledge in extracellular vesicle research. Nat Methods 2017;14:228–32.

[24] Welsh JA, Van Der Pol E, Arkesteijn GJA, et al. MIFlowCyt-EV: a framework for standardized reporting of extracellular vesicle flow cytometry experiments. J Extracell Vesicles 2020.

[25] Welsh JA, Horak P, Wilkinson JS, et al. FCMPASS Software Aids Extracellular Vesicle Light Scatter Standardization. Cytom Part A 2020;97:569–81.

[26] Welsh JA, Jones JC. Small Particle Fluorescence and Light Scatter Calibration Using FCMPASS Software. Curr Protoc Cytom 2020;94.

[27] Welsh JA, van der Pol E, Bettin BA, et al. Towards defining reference materials for measuring extracellular vesicle refractive index, epitope abundance, size and concentration. J Extracell Vesicles 2020;9:1816641.

[28] Exometry B. Rosetta Calibration - Exometry.

[29] van der Pol E, Böing AN, Gool EL, et al. Recent developments in the nomenclature, presence, isolation, detection and clinical impact of extracellular vesicles. J Thromb Haemost 2016;14:48–56.

[30] Paulaitis M, Agarwal K, Nana-Sinkam P. Dynamic Scaling of Exosome Sizes. Langmuir 2018;34:9387–93.

[31] Mallick ER, Witwer KW, Arab T, et al. Characterization of extracellular vesicles and artificial nanoparticles with four orthogonal single-particle analysis platforms. BioRxiv 2020:2020.08.04.237156.

[32] Osteikoetxea X, Sódar B, Németh A, et al. Differential detergent sensitivity of extracellular vesicle subpopulations. Org Biomol Chem 2015;13:9775–82.

[33] Van Der Pol E, Coumans FAW, Sturk A, et al. Refractive index determination of nanoparticles in suspension using nanoparticle tracking analysis. Nano Lett 2014;14:6195–201.

[34] Bachurski D, Schuldner M, Nguyen PH, et al. Extracellular vesicle measurements with nanoparticle tracking analysis–An accuracy and repeatability comparison between NanoSight NS300 and ZetaView. J Extracell Vesicles 2019;8.

[35] Erdbrügger U, Lannigan J. Analytical challenges of extracellular vesicle detection: A comparison of different techniques. Cytom Part A 2016;89:123–34.

[36] Corso G, Heusermann W, Trojer D, et al. Systematic characterization of extracellular vesicles sorting domains and quantification at the single molecule–single vesicle level by fluorescence correlation spectroscopy and single particle imaging. J Extracell Vesicles 2019;8.

[37] García-Santamaría F, Míguez H, Ibisate M, et al. Refractive index properties of calcined silica submicrometer spheres. Langmuir 2002;18:1942–4.

[38] Kasarova SN, Sultanova NG, Ivanov CD, et al. Analysis of the dispersion of optical plastic materials. Opt Mater (Amst) 2007;29:1481–90.

[39] Gardiner C, Shaw M, Hole P, et al. Measurement of refractive index by nanoparticle tracking analysis reveals heterogeneity in extracellular vesicles. J Extracell Vesicles 2014;3.

[40] van der Pol E, Coumans FAW, Grootemaat AE, et al. Particle size distribution of exosomes and microvesicles determined by transmission electron microscopy, flow cytometry, nanoparticle tracking analysis, and resistive pulse sensing. J Thromb Haemost 2014;12:1182–92.

[41] Defante AP, Vreeland WN, Benkstein KD, et al. Using Image Attributes to Assure Accurate Particle Size and Count Using Nanoparticle Tracking Analysis. J Pharm Sci 2018;107:1383–91.

[42] Théry C, Amigorena S, Raposo G, et al. Isolation and Characterization of Exosomes from Cell Culture Supernatants and Biological Fluids. Curr. Protoc. Cell Biol., vol. Chapter 3, Hoboken, NJ, USA: John Wiley & Sons, Inc.; 2006, p. Unit 3.22.

[43] Gardiner C, Di Vizio D, Sahoo S, et al. Techniques used for the isolation and characterization of extracellular vesicles: results of a worldwide survey. J Extracell Vesicles 2016;5:32945.

[44] Royo F, Thery C, Falcon-Perez JM, et al. Methods for Separation and Characterization of Extracellular Vesicles: Results of a Worldwide Survey Performed by the ISEV Rigor and Standardization Subcommittee. Cells 2020;9:1955.

[45] Böing AN, van der Pol E, Grootemaat AE, et al. Single-step isolation of extracellular vesicles by size-exclusion chromatography. J Extracell Vesicles 2014;3:23430.

[46] Linares R, Tan S, Gounou C, et al. High-speed centrifugation induces aggregation of extracellular vesicles. J Extracell Vesicles 2015;4:29509.

[47] Van Deun J, Mestdagh P, Sormunen R, et al. The impact of disparate isolation methods for extracellular vesicles on downstream RNA profiling. J Extracell Vesicles 2014;3:24858.

[48] Arab T, Raffo-Romero A, Van Camp C, et al. Proteomic characterisation of leech microglia extracellular vesicles (EVs): comparison between differential ultracentrifugation and Optiprep™ density gradient isolation. J Extracell Vesicles 2019;8:1603048.

